# Mapping the endosomal proximity proteome reveals Retromer as a hub for RAB GTPase regulation

**DOI:** 10.1101/2024.11.22.622898

**Authors:** Carlos Antón-Plágaro, Kai-en Chen, Qian Guo, Meihan Liu, Ashley J. Evans, Philip A. Lewis, Kate J Heesom, Kevin A. Wilkinson, Brett M. Collins, Peter J. Cullen

**Affiliations:** School of Biochemistry, Faculty of Life Sciences, Biomedical Sciences Building, University of Bristol, Bristol BS8 1TD, UK; The University of Queensland, Institute for Molecular Bioscience, St Lucia, Queensland, 4072, Australia; Bristol Proteomics Facility, School of Biochemistry, Faculty of Life Sciences, Biomedical Sciences Building, University of Bristol, Bristol BS8 1TD, UK; School of Physiology, Pharmacology and Neuroscience, Faculty of Life Sciences, Biomedical Sciences Building, University of Bristol, Bristol BS8 1TD, UK

## Abstract

Endosomal retrieval and recycling of integral cargo proteins is essential for cell, tissue and organism-level development and homeostasis and is orchestrated through a specialised retrieval sub-domain on the endosomal vacuole. However, although sub-domain dysfunction is associated with human disease our appreciation of the molecular details and functional components of the retrieval sub-domain(s) remains poorly described. Here, using comparative proximity proteomics of critical retrieval sub-domain components Retromer and Retriever, their cargo adaptors, and a component of the opposing ESCRT-degradative sub-domain, we provide a data-rich resource that identifies new molecular details of retrieval sub-domain composition and organization, including an unrecognised complexity in the interface of Retromer with RAB GTPases. Combining X-ray crystallography and in silico predictions with extensive biochemical and cellular analysis, we dissect the direct association of Retromer with RAB10 regulators DENND4A, DENND4C, TBC1D1, and TBC1D4, and the RAB35 regulator TBC1D13. Overall, we conclude that the Retromer retrieval sub-domain constitutes a major hub for the regulated switching of selected RAB GTPases and propose that this constitutes a major component of the role of Retromer in neuroprotection.

## INTRODUCTION

Across a variety of cell types three to four-thousand transmembrane proteins enter and are sorted and transported through the human endosomal network (Cullen and Steinberg, 2018). Efficient sorting of proteins including receptors, channels, transporters, polarity cues and adhesion molecules is essential for cellular, tissue and organism-level development and adult homeostasis and adaptation (Cullen and Steinberg, 2018). An increasing body of evidence points to perturbed endosomal transport being a contributing factor and, in some cases a causative factor in human diseases including neurodegenerative and neurological disorders (McMillan et al., 2017; Small et al., 2017; Qureshi et al., 2020), and metabolic syndromes and cardiovascular disease (Gilleron and Zeigerer, 2023). Hijacking the host endosomal network is also an emerging theme in the infectivity of human viral and bacterial pathogens (Simonetti et al., 2023).

On entering the endosomal network transmembrane proteins and their associated proteins and lipids (collectively termed ‘cargos’) are sorted between two fates. A series of multi-protein ESCRT complexes (endosomal sorting complexes required for transport) recognize cargo modified through monoubiquitylation and transport these for degradation within the lysosome (Vietri et al., 2020). Other cargos avoid this degradative fate and undergo retrieval for recycling and further rounds of reuse at organelles that include the cell surface, the biosynthetic and autophagic pathways, and lysosomes and lysosome-related organelles (Cullen and Steinberg, 2018).

Orchestrating endosomal cargo retrieval and recycling are several multi-protein assemblies that include sorting nexin-27 (SNX27)-Retromer and SNX3-Retromer (Harterink et al., 2011; Temkin et al., 2011; Steinberg et al., 2013; Gallon et al., 2014; Clairfeuille et al., 2016; Lucas et al., 2016; Kovtun et al., 2018; McGough et al., 2018; Leneva et al., 2021), SNX17-Retriever and associated Commander super-assembly (Phillips-Krawczak et al., 2015; Mallam and Marcotte, 2017; McNally et al., 2017; Healy et al., 2023; Boesch et al., 2024; Laulumaa et al., 2024), the branched F-actin polymerizing WASH complex and its regulatory MAGE-L2/USP7/TRIM27 (MUST) complex (Derivery et al., 2009; Gomez and Billadeau, 2009; Hao et al., 2015), and the SNX-BAR proteins and endosomal sorting complex required for exit-1 (ESCPE-1) (Carlton et al., 2004, Simonetti et al., 2019; Yong et al., 2020; Yong et al., 2021; Chandra et al., 2022; Simonetti et al., 2022a; Simonetti et al., 2022b).

On the limiting membrane of endosomes, electron and light microscopy has established that core retrieval and recycling complexes localise to specific retrieval sub-domains that while present on the same endosome are physically distinct to ESCRT-demarcated degradative sub-domains (Sachse et al., 2002; Mari et al., 2008; Puthenveedu et al., 2010; Bowman et al., 2016; Varandas et al., 2016; McNally et al., 2017; Norris et al., 2017). Analysis in cells and organisms have established the essential importance of functional retrieval sub-domains for development and homeostasis and disease associated sub-domain dysregulation is associated with human disease (McMillan et al., 2017; Small et al., 2017; Qureshi et al., 2020; Gilleron and Zeigerer, 2023). Defining the organization and function of retrieval sub-domains is central to define the mechanistic basis of endosomal cargo retrieval and recycling.

Although their importance is well established, the precise functional composition of the retrieval sub-domain remains poorly characterised. Here we have employed BioID-based quantitative proteomics to identify the proximity protein environment of the endosomal retrieval sub-domain specifically focusing on Retromer and Retriever and their chief respective cargo adaptors SNX27 and SNX17 and the HRS component of the ESCRT-0 complex. Through strategic placement of the BioID enzyme, we reveal proximity information relating to the known configuration of Retriever within the Commander super-assembly (Healy et al., 2023; Boesch et al., 2024; Laulumaa et al., 2024) and the arch-like architecture of membrane associated Retromer (Kovtun et al., 2018; Leneva et al., 2021). By establishing new molecular components associated with the Retromer retrieval sub-domain we identify a previously unrecognised complexity in Retromer’s interface with RAB GTPases that we structurally and functionally dissect to reveal Retromer as a major endosomal hub for the regulation of a select group of RAB GTPases.

## RESULTS

### Proximity proteomics of endosomal sorting sub-domains

To biochemically probe the organization of the degradative and retrieval sub-domains we employed the proximity-dependent biotin identification methodology (BioID) (Roux et al., 2012; Varnaité and MacNeill, 2016) (**Figure 1A**). Upon addition of membrane permeable biotin, the BioID1 enzyme tagged to a protein of interest (POI) catalyzes the formation of a highly reactive but labile compound, biotinoyl-5’-AMP, which covalently labels lysine residues of neighbouring proteins within a radius of proximity of approximately 10 nm (for context Retromer is approximately 15 - 20 nm in length (Kovtun et al., 2018; Leneva et al., 2021). Processing of resulting biotinylated proteins through cell lysis and streptavidin affinity isolation coupled with isobaric tandem mass tagging (TMT) and nanoscale liquid chromatography joined to tandem mass spectrometry (nano-LC-MS/MS) provides an unbiased view of the local proximity proteome (**Figure 1A**). Several derivations of BioID are available (Sears et al., 2019), but we selected BioID1 because of its long record of use and its low basal activity (Gentzel et al., 2019), ideal for our CRISPR/Cas9 knock out (KO) and chimeric rescue approach outlined below.

**Figure 1.**
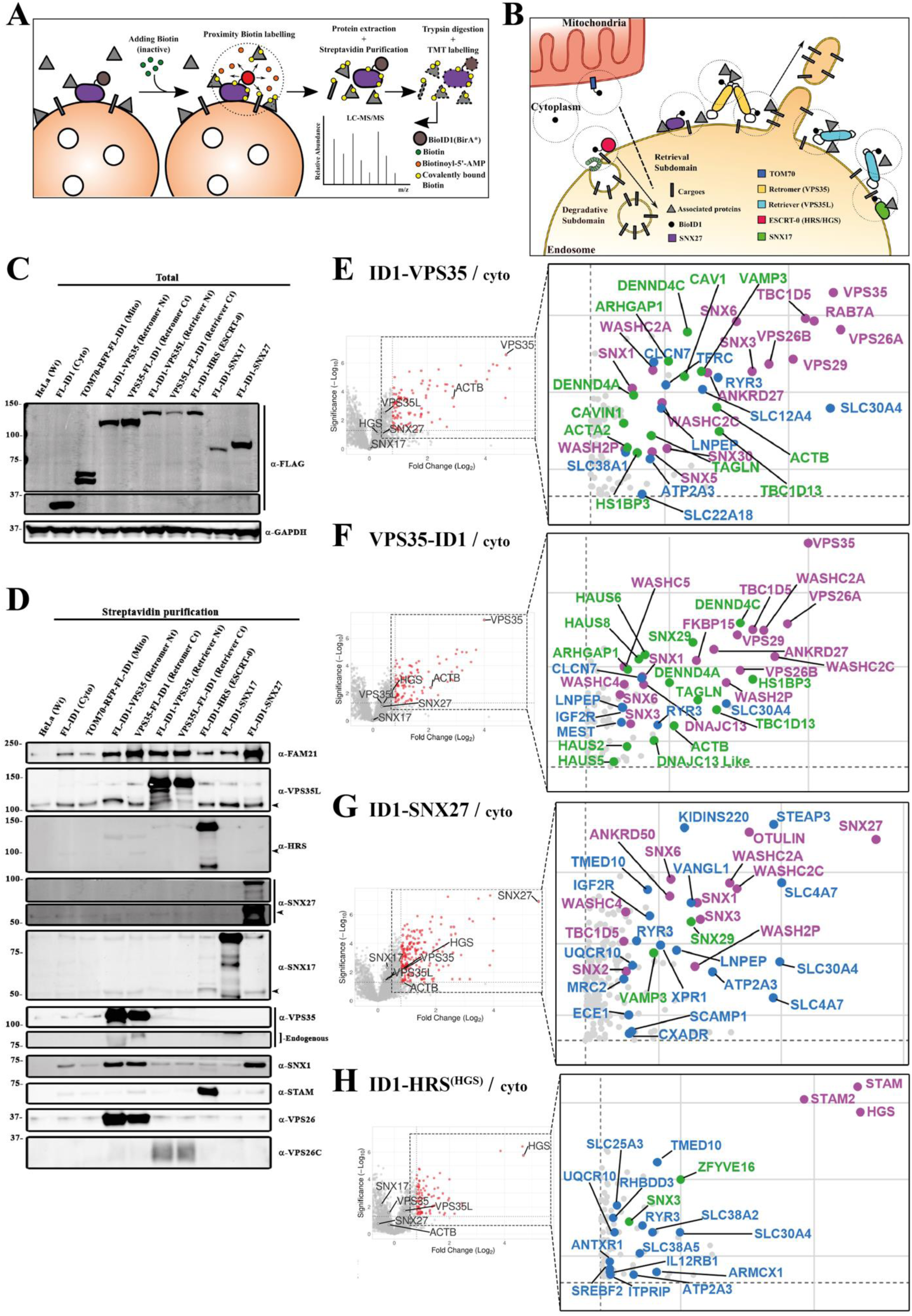
Proximity proteomics of endosomal sorting sub-domains. **(A)** Schematic representation of proximity-dependent biotin identification (BioID) of a protein of interest (POI) over endosomal surface. **(B)** Cartoon depicting endosomal proteins tagged with BioID1, including a soluble version called cytosolic BioID1 and a TOM70-directed cytoplasmic facing mitochondrial external membrane. **(C)** Western blot showing comparison of total protein levels of the BioID1-tagged proteins expressed in the engineered cell lines using antibody against FLAG epitope. **(D)** Western blot comparing biotinylated proteins after 24-hours biotin incubation, lysis and streptavidin purification among the different cell lines. **(E-H)** Volcano plots representing the proteins detected in proximity to the indicated endosomal proteins when compared with the cytosolic BioID1. X axis denotes Fold Change (FC) in Log_2_ scale and Y axis statistical significance in −Log_10_ scale format; for reference 1.3 −Log_10_ is equal to 0.05 p-value. Right insets are magnifications showing only significant proteins (≥ 1.3) with FC ≥ 0.8. In magenta already known interactors, in blue transmembrane proteins, in green new or barely studied interactors. **(E)** BioID1-VPS35. **(F)** VPS35-BioID1. **(G)** BioID1-SNX27. **(H)** BioID1-HRS/HGS.

To biochemically probe the organization of the retrieval sub-domain we focused on: *(i)* the core VPS35 subunit of Retromer for which we generated two constructs BioID1 tagged at either its carboxy or amino-termini (VPS35-BioID1 and BioID1-VPS35 respectively); *(ii)* the core VPS35L subunit of Retriever again either carboxy or amino-termini tagged (VPS35L-BioID1 and BioID1-VPS35L respectively); *(iii)* the major cargo adaptors for Retromer and Retriever, SNX27 and SNX17 respectively, tagged with BioID1 at their amino-termini (BioID1-SNX27 and BioID1-SNX17); and *(iv)* an amino-terminal tagged version of the ESCRT-0 subunit HRS to label the degradative sub-domain (BioID1-HRS). As additional controls for filtering the selectivity of the BioID1 proximity proteomes we generated HeLa lines expressing a soluble version of cytosolic BioID1 and a version of BioID1 directed to the cytoplasmic facing mitochondrial external membrane using the TOM70 signal sequence (TOM70-RFP-BioID1) (**Figure 1B**). In all cases, coupling to BioID1 included a short flexible linker (GGGGGGKGA) and a FLAG epitope organised with respect to the protein of interest (POI) as follows; POI-Linker-FLAG-BioID1 or FLAG-BioID1-Linker-POI.

To express these constructs in HeLa cells we adopted a two-step approach. We first used CRISPR/Cas9 KO to generate individual clonal HeLa lines for all gene alleles. Into the relevant lines we used lentiviruses to transduce the corresponding BioID1 tagged chimera, and by means of viral titration and puromycin selection we isolated populations expressing as near to endogenous levels of the chimera as possible (**Figure 1C**). We extensively validated the localization and function of each resulting cell population. Endosomal localization was confirmed by immuno-fluorescence using specific antibodies or where suitable antibodies were not available through anti-FLAG labelling (***Supplementary Figure 1***). Functionality was defined through the ability of each chimera to rescue well-established missorting phenotypes of endosomal cargo proteins observed upon KO of SNX27-Retromer and SNX17-Retriever complexes (Steinberg et al., 2013; McNally et al., 2017). All chimeric versions of VPS35 and SNX27 rescued the lysosomal missorting of the GLUT1 glucose transporter and promoted its recycling to the cell surface (***Supplementary Figure 2A and 2B***). Similarly, SNX17 and the VPS35L chimeras rescued the lysosomal missorting of α5β1-integrin and promoted its cell surface recycling (***Supplementary Figure 2C and 2D***). The chimeric version of HRS was able to rescue the loss of expression of the other component of the ESCRT-0 complex STAM1/STAM2 and rescued the morphology of the EEA1 and SNX1-positive endosomes, consistent with its integration into a functional complex (***Supplementary Figure 2E-2G)*.**

To explore the robustness of the proximity labelling we performed an initial biased identification of the biotinylated proteins using quantitative western analysis of target proteins. Although 12 hours incubation with biotin was enough to label some interactors, we chose to perform all experiments using 24 hours incubation to maximize the detection of weak and low abundance proximity proteins, and the potential to identify cargo proteins transiting through the sub-domains (***Supplementary Figure 2H***). Importantly, we tested whether the actual biotin-labelling process had any detectable effect on Retromer and Retriever-dependent cargo sorting. After 24 hours of incubation with 50 mM biotin, we did not detect significant changes in the trafficking of GLUT1 or α5β1-integrin in any of the HeLa lines expressing the chimeric versions (***Supplementary Figure 2A-D***). This suggests that the BioID1 protocol does not induce a detectable perturbation in endosomal sorting through a build-up of inhibitory biotin on the functional machinery within the sorting sub-domains.

Across the different BioID1 chimeric HeLa cell lines the level of each biotinylated protein was comparable when blotted using the same FLAG antibody, being heterogeneous but always lower than the cytosolic BioID1 (**Figure 1C**). Parallel comparison of the biotinylated proteins after 24 hours incubation established minimal labelling of SNX27-Retromer, SNX17-Retriever, and HRS by parental HeLa cells lacking any BioID1 expression, and HeLa cells expressing cytoplasmic BioID1 or TOM70-RFP-BioID1 (**Figure 1D**). For Retromer, BioID1-VPS35 and VPS35-BioID1 revealed strong labelling of endogenous VPS26 and FAM21, a component of the WASH complex that directly associates with VPS35 (Gomez and Billadeau, 2009; Harbour et al., 2012; Jia et al., 2012; Guo et al., 2023). Labelling was also detected for SNX1 a component of the ESCPE-1 complex (Simonetti et al., 2019) but unexpectedly not SNX27 (Steinberg et al., 2013; Gallon et al., 2014). For Retriever, VPS35L-BioID1 and BioID1-VPS35L labelled endogenous VPS26C and FAM21, consistent with the known indirect association of Retriever with the WASH complex (McNally et al., 2017). BioID1-SNX17 showed limited labelling for the blotted proteins. In contrast, BioID1-SNX27 showed robust labelling of FAM21 and SNX1 entirely consistent with its known direct interaction with these proteins (Steinberg et al., 2013; Yong et al., 2021; Chandra et al., 2022; Simonetti et al., 2022; Guo et al., 2023). Finally, BioID1-HRS labelled endogenous STAM1 with minimal detectable binding to any proteins from Retromer, Retriever, and the ESCPE-1 and WASH complexes. These data provide proof-of-principle that the experimental design is suitable for providing biochemical insight into the organization of the endosomal sorting sub-domains.

### Unbiased quantitative proximity proteomics of endosomal sorting sub-domains

To unbiasedly identify the proximity proteomes, we performed 10-plex TMT-based quantitative proteomics across 6-independent experimental repeats comparing the protein abundances between BioID1-VPS35, VPS35-BioID1, BioID1-VPS35L, VPS35L-BioID1, BioID1-SNX17, BioID1-SNX27, BioID1-HRS and the parental, cytoplasmic BioID1 and TOM70-RFP-BioID1 controls. Resulting data were compared between the individual endosomal BioID1 chimera and the cytosolic BioID1 in volcano plots of the Fold Change (Log_2_) versus statistical significance (-Log_10_) (**Figures 2E-H and 3A-C**).

**Figure 2.**
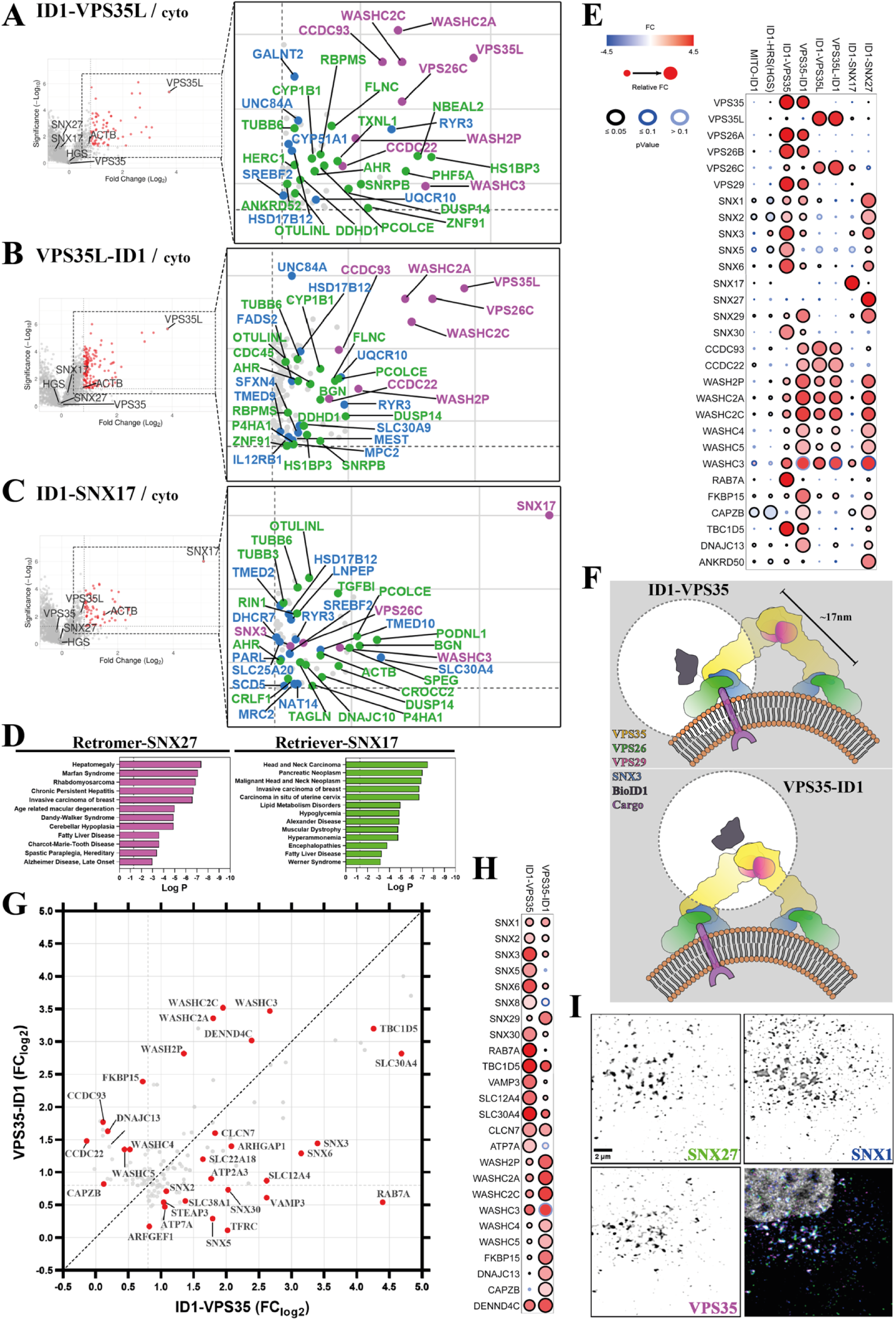
Proximity proteomes of Retriever (VPS35L) and SNX17, and comparative gene ontology (GO) comparison among all proximity proteomes. **(A-C)** Volcano plots representing the proteins detected in proximity to the indicated endosomal proteins when compared with the cytosolic BioID1. X axis denotes Fold Change (FC) in Log_2_ scale and Y axis statistical significance in −Log_10_ scale format; for reference 1.3 −Log_10_ is equal to 0.05 p-value. Right insets are magnifications showing only significant proteins with FC ≥ 0.8. In magenta already known interactors, in blue transmembrane proteins, in green new or barely studied interactors. **(A)** BioID1-VPS35L. **(B)** VPS35L-BioID1. **(C)** BioID1-SNX17. **(D)** Selected disease associated GO terms and its significance (-Log_10_) for combined data of SNX27-Retromer and SNX17-Retriever proximity proteomes. **(E)** Dot plot showing the comparison of FC (Log_2_) and significance of selected endosomal proteins and Mito-BioID1. Filling colours represent the FC value, edge colours the p-value and the size symbolize the relative FC. **(F)** Cartoon depicting the expected localization of the BioID1 biotin ligase, in black, along the arch-like conformation of Retromer over endosomal surface. Edge-dashed circles represent the 10 nm-radius spheres where BioID1 can biotinylate proteins. Sizes are based on Retromer arch structure PDB:7BLN. **(G)** Scatter plot showing the FC values (Log_2_) for detected proteins in both Retromer proximity proteomes; FC from BioID1-VPS35 in X axis and FC from VPS35-BioID1 in Y axis. The farther from the central dashed line the protein is, the greater the preferential proximity labelling to a specific BioID1 version. **(H)** Dot plot showing the comparison of FC (Log_2_) and significance of selected endosomal proteins. **(I)** Colocalization of endogenous Retromer (VPS35), SNX27 and SNX1 in HeLa cells. Scale bar - 2 μm.

As expected, both BioID1-VPS35 and VPS35-BioID1 labelled the other Retromer subunits VPS29, VPS26A and VPS26B and several previously identified Retromer accessory proteins including SNX3, RAB7A, TBC1D5, ANKRD27, DENND4A and DENND4C, FKBP15, and the WASHC2A (FAM21A/B), WASHC2C (FAM21C), WASHC5 (strumpellin), WASHC4 (SWIP) and WASH2P (WASH) subunits of the WASH complex (Rojas et al., 2008; Gomez and Billadeau, 2009; Seaman et al., 2009; Harterink et al., 2011; Harbour et al., 2012; Jia et al., 2012; Hesketh et al., 2014; McGough et al., 2014; McMillan et al., 2016) (**Figure 1E and 1F**).

For BioID1-VPS35L and VPS35L-BioID1, the Retriever subunit VPS26C was identified along with the coiled coil proteins CCDC22 and CCDC93 which form the backbone of the CCC complex that associates with Retriever to form, alongside DENND10, the Commander super-assembly (Phillips-Krawczak et al., 2015; McNally et al., 2017; Healy et al., 2023; Boesch et al., 2024; Laulumaa et al., 2024) (**Figure 2A and 2B**). The identification of WASHC2C (FAM21C), WASHC2A (FAM21A/B), WASHC3 (SWIP) and WASH2P (WASH) is consistent with the known association of the CCC complex to the WASH complex (Phillips-Krawczak et al., 2015) (**Figure 2A and 2B**). Finally, ID1-HRS/HGS showed robust labelling of the other ESCRT-0 subunits STAM1 and STAM2, but there was no significant labelling of Retromer and associated accessory proteins (the exception being SNX3), Retriever or the CCC and WASH complexes (**Figure 1H**). Together these data provide an unbiased validation of the experimental design and reveal little proximity overlap between the molecular components of the ESCRT-0 demarcated degradative sub-domain and the retrieval sub-domain defined by VPS35 and VPS35L.

For the Retromer cargo adaptor SNX27, ID1-SNX27 labelled a wide range of known interactors including the WASH subunits WASHC2A (FAM21A/B), WASHC2C (FAM21C), WASHC4 (SWIP) and WASH2P (WASH), the ESCPE-1 subunits SNX1, SNX2 and SNX6 (Simonetti et al., 2019; Yong et al., 2021; Chandra et al., 2022; Simonetti et al., 2022; Guo et al., 2023), ANKRD50 (Kvainickas et al., 2017) and Otulin (Stangl et al., 2019) (**Figure 1G**). Multiple transmembrane proteins were also labelled many of which contain PDZ binding motifs and rely on SNX27 for their endosomal retrieval and recycling; examples included KIDINS220, STEAP3, SLC4A7 and VANGL1 (Steinberg et al., 2013; Clairfeuille et al., 2016) (**Figure 1G**). Surprisingly, while BioID1-SNX27 labelled the core Retromer subunit VPS35 it failed to reach the cut-off for log_2_ fold change (≥ 0.8) and significance (**Figure 1G**), and similarly BioID1-VPS35 or VPS35-BioID1 while labelling SNX27 failed to reach the same cut-off parameters (**Figure 1E and 1F**). Given the direct nature of the PDZ domain of SNX27 in binding to VPS26A and VPS26B and the importance of this coupling for Retromer-mediated cargo sorting (Gallon et al., 2014), the lack of significantly enriched labelling may result from a technical limitation or reflect a highly dynamic, transient association between cargo-loaded SNX27 and the Retromer demarcated sub-domain during the handover of captured cargo (Simonetti et al., 2022). BioID1-SNX17 on the other hand labelled the VPS26C subunit of Retriever consistent with the mechanism of SNX17 binding to this heterotrimer (McNally et al., 2017; Butkovic et al., 2024; Singla et al., 2024) (labelling of VPS35L fell just below the log_2_ fold change cut-off) (**Figure 2C**).

Unbiased analysis of all the significant protein hits with FC ≥ 0.8 (log_2_) using metascape.org platform (Zhou et al., 2019), revealed top enriched ontology terms for categories including Cellular Component, Biological Processes and CORUM (***Supplementary Figure 3***). With the expected exception of Mito-ID1, all protein lists showed significant enrichment for the GO term endosomal transport and additional terms including ‘endocytic recycling’, ‘regulation of transport’, ‘Retromer complex, Retromer complex binding and Retromer tubulation complex’, ‘CCDC22-CCDC93-C16orf62-FAM21-WASH complex’, ‘WASH complex’, and ‘regulation of actin filament-based process’. From analysis of Disease Processes, the data set revealed a significant enrichment of the GO terms associated with a range of neurological conditions, metabolic syndromes and cancers consistent with the emerging functional role of the endosomal retrieval sub-domain in neuroprotection, metabolic regulation and growth factor signalling (**Figure 2D**, ***Supplementary Figure 3***). Overall, these data further validate the experimental design and reveal the global strength of the acquired proximity proteomes.

### Quantitative comparison among endosomal sub-domains

As our 10-Plex TMT-based experimental design allowed all proximity proteomes to be collected within the same proteomic run, we used the platform prohits-viz.org (Knight et al., 2017) and specifically the dot plot option to better visualize the quantitative enrichment and statistical significance of each identified protein across all data sets (***Supplementary Figure 4***). For the Retromer VPS26A/B:VPS35:VPS29 and Retriever VPS26C:VPS35L:VPS29 complexes this allowed the clear visualization that ID1-VPS35 and VPS35-ID1 displayed a significant enrichment of VPS26A, VPS26B and VPS29, an enrichment that was not detected through BioID1 tagging of VPS35L, HRS (ESCRT-0 subunit) or the MITO-ID1 (**Figure 2E**). The same was also true for VPS35L-ID1 and ID1-VPS35L where VPS26C was clearly detected but only moderately detected by ID1-VPS35. The lack of VPS29 enrichment in either VPS35L proximity proteomes may reflect a technical caveat or arise from a restricted capacity for biotinylation due to the distinct mechanism for VPS29 association within the Retriever assembly (Healy et al., 2023; Boesch et al., 2024; Laulumaa et al., 2024). These data point to a level of spatial segregation between Retromer and Retriever within the retrieval sub-domain that, as far as we are aware, has not been resolved by imaging. That said, the membrane distal VPS35-ID1 but not the membrane proximal ID1-VPS35 (see below for more detailed discussion) catalyzed labelling of specific subunits of the Retriever associated CCC complex, namely CCDC22 and CCDC93, suggestive of a proximity between Retromer and Retriever containing Commander super-assembly. Building on this, the common labelling of the WASH complex by both Retromer and Retriever (**Figure 2E**) is entirely consistent with their known biochemical association and physical co-localisation to the retrieval sub-domain and the central role of actin polymerization and actin turnover in Retromer- and Retriever-dependent cargo sorting (Gomez and Billadeau, 2009; Harbour et al., 2010; Gomez et al., 2012; Phillips-Krawczak et al., 2015; Bartuzi et al., 2016; McNally et al., 2017). Indeed, β-actin was detected by Retromer and Retriever BioID1s, but not in BioID1-HRS, consistent with the known segregation of branched filamentous actin with the retrieval sub-domain (***Supplementary Figure 4***).

### Proximity proteomics provides biochemical insight into Retromer’s organisation in the retrieval sub-domain

Cryo-ET analysis of membrane associated Retromer has established a polarized organization where the amino-terminal region of VPS35 associates with VPS26A/B proximal to the membrane surface and the membrane distal carboxy-terminal region of VPS35 dimerizes with a neighbouring Retromer to form an arch-like structure (**Figure 2F**; ***Supplementary Figure 4B***) (Lucas et al., 2016; Kovtun et al, 2018; Leneva et al, 2021). By including two BioID1 versions of VPS35, we sought to provide *in cellulo* biochemical insight into the organization of Retromer relative to the endosomal surface, reasoning that the biotin ligase in ID1-VPS35 should be relatively close to the endosomal membrane (**Figure 2F**, **upper panel**), while in the VPS35-ID1 version it should be towards the top of the arches, more distal to the membrane (**Figure 2F**, **lower panel**). For those identified proteins with FC ≥ 0.8 we plotted ID1-VPS35 versus VPS35-ID1 to observe any quantitative difference in labelling efficiency across the proximity proteomes (**Figure 2G**). This revealed that the membrane proximal ID1-VPS35 preferentially labelled several transmembrane proteins including well-established Retromer cargo proteins such as STEAP3, TFRC and ATP7A (**Figure 2G**) (Steinberg et al., 2013). At the mechanistic level, ID1-VPS35 preferentially labelled the membrane anchored RAB7 GTPase, which plays a central role in the endosomal recruitment of Retromer through binding to VPS35 (Rojas et al., 2008; Seaman et al., 2009), and the phosphatidylinositol 3-monophosphate (PI(3)P) binding sorting nexin SNX3, which by associating with the VPS26A/B:VPS35 interface assists in the endosomal association of Retromer and the sequence-dependent recognition of cargo undergoing SNX3-Retromer mediated endosomal sorting (**Figure 2G**) (Strochlic et al., 2007; Harterink et al., 2011; Chen et al., 2013; Lucas et al., 2016; McGough et al., 2018; Leneva et al., 2021). Additional membrane binding sorting nexins of the ESCPE-1 complex (Simonetti et al., 2019; Lopez-Robles et al., 2023), namely SNX1, SNX2, SNX5 and SNX6, were also preferentially labelled by ID1-VPS35 (**Figure 2G and 2H**). In addition, and consistent with the colocalization of endogenous VPS35, SNX27 and SNX1 (**Figure 2I**, ***Supplementary Figure 4D***), ID1-SNX27 also labelled ESCPE-1 subunits (**Figure 2E**) supporting the recently proposed cargo handover model for SNX27-Retromer mediated ESCPE-1-dependent cargo sorting (Simonetti et al., 2022). Here the SNX1 and SNX2 subunits of ESCPE-1 bind directly to the FERM domain of SNX27 to couple sequence-dependent cargo recognition with the biogenesis of tubular transport carriers for the promotion of plasma membrane recycling (Yong et al., 2021; Chandra et al., 2022; Simonetti et al., 2022). Indeed, the relative FC of the ID1-SNX27 labelling of SNX1 and SNX2 relative to SNX5 and SNX6 is entirely consistent with this mechanism of coupling (**Figure 2E**).

In contrast to the labelling of membrane proximal proteins by ID1-VPS35, all subunits of the WASH complex were preferentially detected by the membrane distal VPS35-ID1 (**Figure 2G and 2H**). This is entirely consistent with the mechanism of WASH complex association where acidic-Asp-Leu-Phe (aDLF) motifs in the FAM21 subunit bind to VPS29 and two sites towards the carboxy-terminal membrane distal region of VPS35 (Guo et al., 2023). Two additional regulators of endosomal actin dynamics the FK506-binding protein-15 (FKBP15) (Viklund et al., 2009; Harbour et al., 2012; Nooh and Bahouth, 2017) and the actin binding and capping proteins CAPZA2 and CAPZB (Wear and Cooper, 2004; Wang et al., 2021) were also preferentially labelled by VPS35-ID1 (**Figure 2G and 2H**), as was DNAJC13 (also known as RME-8) a protein that by catalyzing the removal of encroaching HRS from the degradative sub-domain serves to regulate the separation of degradative and retrieval sub-domains (**Figure 2G and 2H**) (Shi et al., 2009; Popoff et al., 2009; Freeman et al., 2014, Norris et al., 2017; Norris et al., 2022). Together, these data establish biochemical signatures for the inner membrane proximal and outer membrane distal layers of key accessory proteins of the Retromer coat complex.

In performing a similar analysis with ID1-VPS35L and VPS35L-ID1 we failed to detect such clear segregation in the proximity profiles of Retriever accessory proteins (**Figure 2E**; ***Supplementary Figure 4***). One likely reason is that Retriever (and the larger Commander assembly) may adopt a completely different membrane-proximal orientation to Retromer. It may also reflect the distinct organization of the Retriever heterotrimer where the amino-and carboxy-termini are less spatially segregated due to its more compact structure (Healy et al., 2023; Boesch et al., 2024; Laulumaa et al., 2024) (***Supplementary Figure 4C***). Although VPS35L BioID1 constructs robustly labelled CCDC22 and CCDC93, we did not detect significant enrichment of any COMMD proteins that form the core decameric ring of the CCC complex. This provides *in cellulo* evidence in support of the proposed structural organization of the Commander super-assembly (Healy et al., 2023; Boesch et al., 2024; Laulumaa et al., 2024).

### Retromer is a hub for GTPase regulation

Having established the robustness of the experimental design and methodology by focusing on known features of Retromer and Retriever we next probed for new molecular insight into retrieval sub-domain organisation. Taking those proteins identified across all proximity proteins (396 proteins with FC ≥ 0.8 (log_2_)) we manually sub-classified the data based on functional terms (**Figure 3A**). Within these clusters we searched for novel proteins whose enrichment and statistical significance profiles displayed a heavy bias towards detection by VPS35-ID1 and/or ID1-VPS35 (proteins highlighted by red arrows in **Figure 3A**). This identified several potential new components of the Retromer sub-domain including the endosome-to-TGN SNARE VAMP3, sorting nexin-29 (SNX29), the caveolae associated caveolin-1 (CAV1) and CAVIN1, subunits of the octameric augmin complex (HAUS2, HAUS5, HAUS6, HAUS8), which interacts with the γ-tubulin ring complex (γTuRC) to stabilize pre-existing microtubules and facilitate microtubule branching during chromosomal segregation and neuronal migration, development, and polarization (Uehara et al., 2009; Cunha-Ferreira et al., 2018; Gabel et al., 2022; Zhang et al., 2022; Zupa et al., 2022; Travis et al., 2023), and the late endosomal 2Cl^-^/H^+^ exchanger CLCN7 – dysfunction in this exchanger leads to a lysosomal storage disease and neurodegeneration in humans (Kasper et al., 2005). VPS35-BioID1 and BioID1-VPS35 proximity labelling followed by targeted western analysis confirmed the selective labelling of VAMP3, CAV1 and CAVIN1, and subunits of the augmin complex within the Retromer sub-domain, suggestive of these proteins and complexes associating with and/or transiently traversing through this sub-domain (**Figure 3B and 3C**).

**Figure 3.**
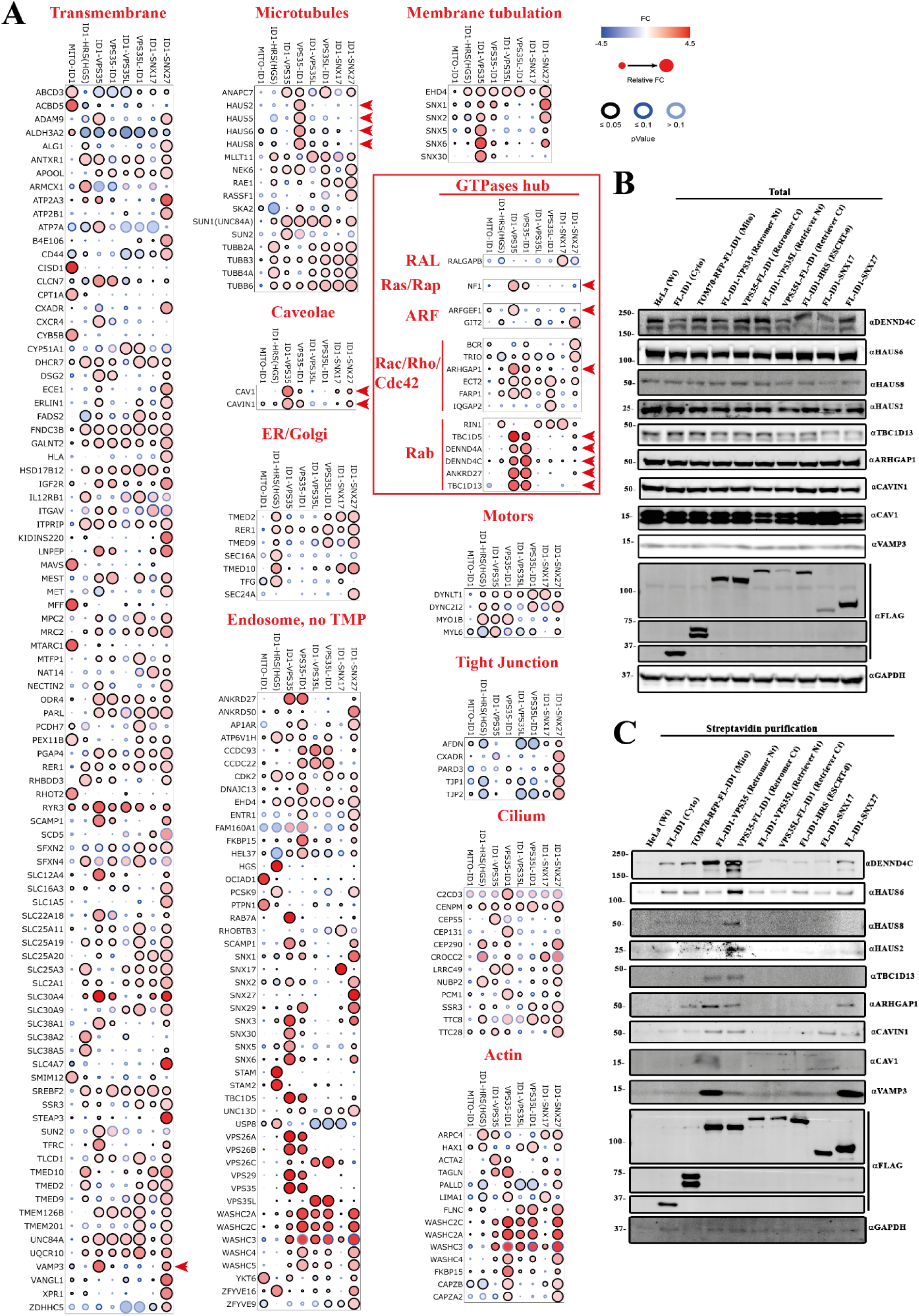
Functional clustering comparing the proximity proteomes. **(A)** Dot plots organized by function, pathway or topology. Red arrow heads highlight proteins displaying a heavy bias towards detection by VPS35-ID1 and/or ID1-VPS35. **(B)** Comparison of total protein levels among all the BioID1 cell lines using antibodies for potential new Retromer interactors. **(C)** Comparison of the biotinylated proteins after 24-hours biotin incubation, lysis and streptavidin purification among the different cell lines using antibodies targeting new Retromer proximity proteins.

Also present in the Retromer sub-domain were a variety of GTPase-activating proteins (GAPs) and guanine nucleotide exchange factors (GEFs). These included known Retromer accessory proteins, the RAB7 GAP TBC1D5 (Seaman et al., 2009; Jia et al., 2016; Jimenez-Orgaz et al., 2018; Seaman et al., 2018; Borg Distefano et al., 2018; Kvainickas et al., 2019), the RAB21/RAB32/RAB38 GEF VARP (*a.k.a.* ANKRD27) (Hesketh et al., 2014; McGough et al., 2014; Crawley-Snowdon et al., 2020), and the RAB10 GEFs DENND4A and DENND4C (Yoshimura et al., 2010; Sano et al., 2011). Activation of RAB10 is heavily implicated in general endosomal cargo recycling (Chen et al., 2006; Shi et al., 2010; Shi et al., 2012; Wang et al., 2016; Liu et al., 2018; Etoh and Fukuda, 2019; Brumfield et al., 2021; Khan et al., 2022; Kluss et al., 2022) and one of its effector proteins TBC1D13, a functional RAB35 GAP was also labelled within the Retromer proximity proteome (Davey et al., 2012) (**Figure 3A**). RAB35 itself regulates endosomal exit for cell surface recycling (Kouranti et al., 2006; Sato et al., 2008; Allaire et al., 2010; Chesneau et al, 2012; Kobayashi and Fukuda, 2012), in part through controlling localized actin dynamics and phosphoinositide metabolism (Zhang et al., 2009; Rahajeng et al., 2012). Also labelled within the Retromer proximity proteome was the Cdc42 GAP ARHGAP1 (*a.k.a.* cdc42GAP), and two proteins selectively labelled by ID1-VPS35 over VPS35-ID1, the Ras GAP neurofibromin (NF1) and ARFGEF1 (*a.k.a.* BIG1), a GEF linked with regulation of the Golgi localized ARF1 and ARF3 GTPases (**Figure 3A**).

VPS35 BioID1 proximity labelling followed by targeted western analysis confirmed the selectivity of labelling for these proteins within the Retromer sub-domain (**Figure 3B and 3C**). To probe for the potential direct association of these GEFs and GAPs with Retromer we turned to AlphaFold2 modelling. Consistent with published structural data, AlphaFold2 predicted the association of a conserved surface-exposed hydrophobic cavity on VPS29 (defined by Leu152) with Pro-Leu (PL) motifs from TBC1D5 (^141^PL^142^) and VARP (^713^PL^714^) and for TBC1D5 the association of an additional ^283^Ile-Pro-Phe^285^ hydrophobic motif with VPS35 (***Supplementary Figure 5***) (Jia et al., 2016; Crawley-Snowdon et al., 2020). While AlphaFold2 failed to predict confident models for ascribing ARHGAP1, ARFGEF1 or NF1 binding to Retromer (data not shown), highly confident models were derived for Retromer association with DENND4A and DENND4C (***Supplementary Figure 6***).

The three DENND4 homologues in humans DENND4A, 4B and 4C all have similar domain structures consisting of an amino-terminal MABP domain (MVB12-associated β-prism domain), DENN domain (consisting of the three structural modules uDENN, cDENN and dDENN), a long extended unstructured linker sequence, and a carboxy-terminal domain that has a mixed α/β globular structure of unknown function (***Supplementary Figure 7***). The carboxy-terminal domain is predicted to fold into a globular structure that forms a tight intramolecular association with the amino-terminal MABP-DENN domain module. Searches using Foldseek (van Kempen et al., 2023) suggest that it is a unique structure found only in the DENND4 family members. Both DENND4A and DENND4C are predicted to interact with Retromer using identical mechanisms, and we discuss DENND4C for simplicity. AlphaFold2 modelling predicts that a conserved PL motif (^1203^PL^1204^) associates with the Leu152 containing hydrophobic cavity of VPS29 (**Figure 4A**). Consistent with these models, GFP-nanotrap immuno-isolation of GFP-VPS29 revealed clear association with endogenous DENND4C, an association that like TBC1D5 was largely dependent on the conserved surface-exposed hydrophobic cavity of VPS29 as shown by VPS29 mutations L152E and V174D (**Figure 4B and 4C**). ITC analysis of a PL-containing synthetic peptide from DENND4C established direct binding to recombinant VPS29 with an affinity (*K*_d_) of 15.4 μM, that was blocked by mutation of the PL motif or by the addition of a competing VPS29-binding macrocyclic peptide RT-D1 (**Figure 4D**, and **Table 1**) (Chen et al., 2021). We determined the crystal structure of VPS29 in complex with the DENND4C ^1203^PL^1204^-containing peptide which completely validated the mode of association (**Figure 4E – 4G**). Furthermore, the crystal structure confirmed the prediction that DENND4C engages VPS29 through both the core PL motif as well as an extended upstream sequence that forms a β-strand extension to the VPS29 structure. AlphaFold2 analysis also predicted a similar mechanism of binding to DENND4A that we confirmed by immuno-isolation (**Figure 4B and 4C**). Despite their presence in the BioID proximity proteomes, no association was detected for ARHGAP1, ARFGEF1 or NF1 in co-immunoprecipitation and Western blot analyses (***Supplementary Figure 6H***).

**Figure 4.**
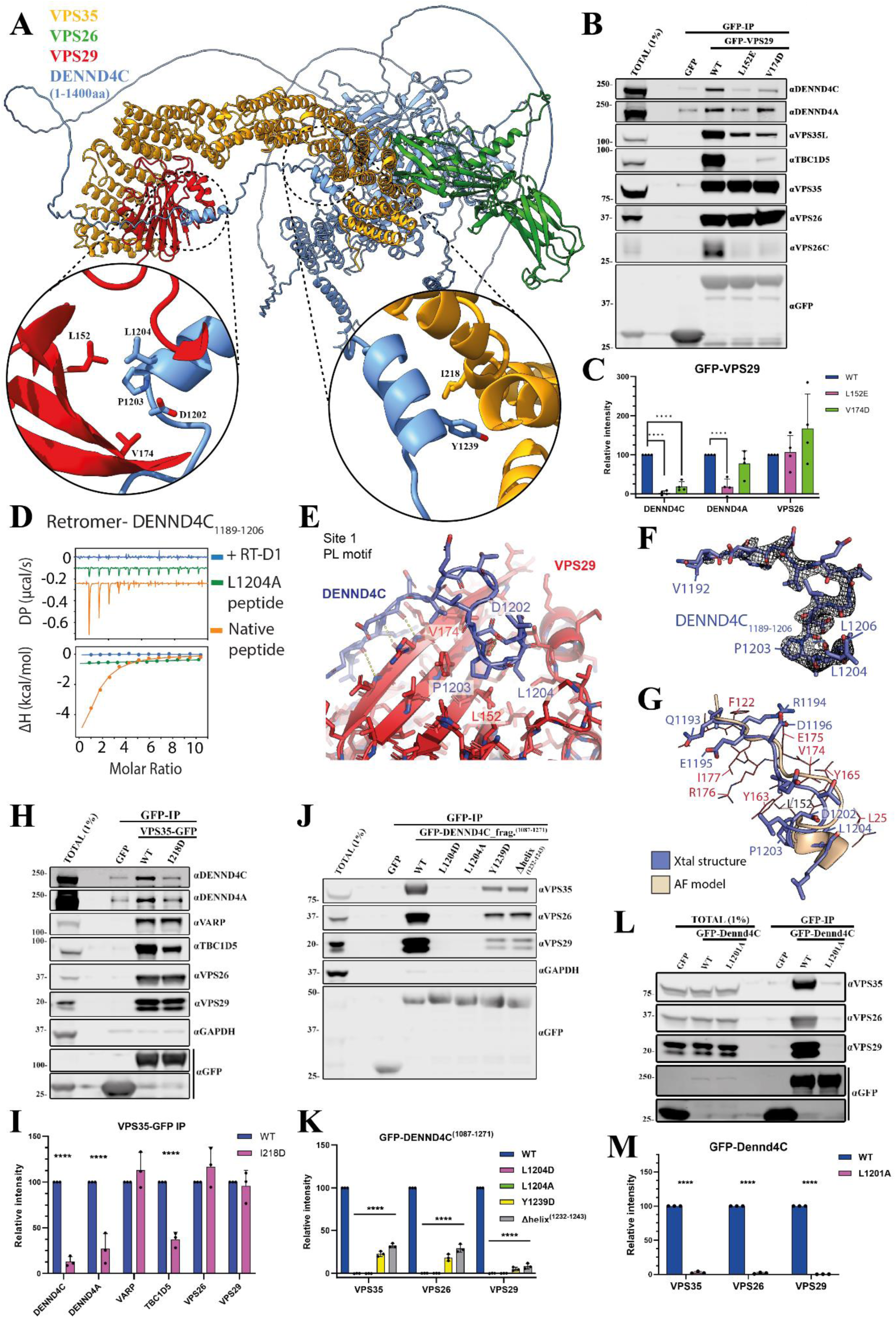
Molecular details of DENND4A and DENND4C binding to Retromer. **(A)** AlphaFold2 prediction of DENND4C (1-1400 aa) interacting with Retromer (rank 1 prediction). Left inset, primary binding site consisting of a conserved PL motif (DENND4C ^1203^PL^1204^) interacting with the Leu152 containing hydrophobic cavity of VPS29. Right inset, second binding site predicted between a short DENND4C α-helical stretch (residues 1232-1243) with α-helices towards the amino-termini of VPS35. **(B, C)** GFP based co-immunoprecipitation (co-IP) and quantification of GFP-VPS29 wild-type (WT) and hydrophobic pocket mutants after transient transfection in HEK293T cells. Quantitation and statistical analysis of relative band intensity for the indicated proteins normalized to GFP band intensity. n = 3 independent experiments. 1-way ANOVA with Dunnett’s multiple comparison test, data presented as mean values relative to WT and error bars represent standard deviation (SD). **(D)** ITC of DENND4C peptides (1189-1206 aa) binding to Retromer in the absence and presence of competing cyclic peptide RT-D1. **(E)** Crystal structure of VPS29 in complex with the DENND4C PL motif-containing peptide^1189-1206^ (PDB:8VOD). **(F)** Ribbon representation of the DENND4C PL-motif containing peptide^1189-1206^. The electron density shown corresponds to a simulated-annealing OMIT Fo - Fc map contoured at 3σ. **(G)** Overlay of the DENND4C peptide structures (slate blue) derived from the crystal structure and from AlphaFold2 prediction (wheat). **(H, I)** GFP based co-IP of VPS35-GFP WT and I218D mutant after transient transfection in HEK293T cells. Quantitation and statistical analysis of relative band intensity normalized to GFP band intensity. n = 3 independent experiments. T-test analysis, data presented as mean values relative to WT and error bars represent SD. **(J, K)** GFP based co-IP of GFP-DENND4C fragment^1087-1271^, WT and mutants after transient transfection in HEK293T cells. Quantitation and statistical analysis of relative band intensity for the indicated proteins normalized to GFP band intensity. n = 3 independent experiments. 1-way ANOVA with Dunnett’s multiple comparison test, data presented as mean values relative to WT and error bars represent SD. **(L, M)** GFP based co-IP after cell lysis of HEK293T cells stably expressing GFP or GFP-Dennd4C full-length from mouse, WT and L1201A mutant; equivalent to human L1204A. Quantitation and statistical analysis of relative band intensity for the indicated proteins normalized to GFP band intensity. n = 3 independent experiments. T-test analysis, data presented as mean values relative to WT and error bars represent SD.

**Table 1.** Thermodynamic parameters for the binding of Retromer and DENND4C by ITC.

| | $K_d$<br>(μM) | $\Delta H$ (kcal/mol) | $\Delta G$<br>(kcal/mol) | $-T\Delta S$<br>(kcal/mol) |
| --- | --- | --- | --- | --- |
| **hRetromer** |  |  |  |  |
| hDENND4C<sub>1189-1206</sub> | 15.4 ± 0.85 | -10.1 ± 1.41 | -6.6 ± 0.04 | -3.5 ± 1.44 |
| hDENND4C<sub>1189-1206</sub> L1204A |  | No binding detected |  |  |
| hDENND4C<sub>1189-1206</sub> + RT-D1 |  | No binding detected |  |  |

While the observed binding of DENND4C to VPS29 is consistent with previous related crystal structures, a second binding site is also predicted between a short DENND4C α-helical stretch (residues 1232-1243) with α-helices towards the amino-termini of VPS35 (**Figure 4A**; ***Supplementary Figure 6***). Consistent with this AlphaFold2 model, immuno-isolation of VPS35-GFP showed association with endogenous DENND4C that was partially reduced upon mutation of residue Ile218 at the amino-termini of VPS35 (**Figure 4H and 4I**). This suggests that DENND4C (and DENND4A) engages Retromer at two distinct sites: a primary site on VPS29, and a secondary site towards the amino-terminal region of VPS35. Consistent with this, immuno-isolation of GFP-tagged DENND4C (a fragment encoding residues 1087-1271) confirmed binding to endogenous Retromer (**Figure 4J and 4K**). This was almost completely lost in mutants targeting the PL motif, Leu1204 (DENND4C(L1204A) and -(L1204D)), and significantly reduced by targeting the secondary site through deletion of the short α-helix (residues 1232-1243) or targeted mutagenesis of Tyr1239 (**Figure 4J and 4K**). ConSurf analysis of amino acid sequence conservation revealed a high degree of evolutionary conservation for both interacting interfaces (***Supplementary Figure 6F***) (Ashkenazy et al., 2016).

Confirming the *in cellulo* relevance of the Retromer interaction with DENND4A and DENND4C, the colocalization of endogenous DENND4C with the endosomal marker SNX1 was partially reduced in a VPS35 knock out HeLa cell line (***Supplementary Figure 8A and 8B***), and while full-length GFP-Dennd4C associated with Retromer decorated endosomes this association was lost with a GFP-Dennd4C(L1201A) mutant that lacked the ability to bind to Retromer, this mutant targets the equivalent Leu1204 PL motif residue in human DENND4C (**Figure 4L and 4M, 5A and 5B**). Double DENND4A and DENND4C knock out in HeLa cells (DENND4A+C KO) revealed a partial defect in the steady-state cell surface levels of GLUT1 (**Figure 5C and *Supplementary Figure 8C***) ***and 8E***). Rescue of the DENND4C+A KO cells with GFP-Dennd4C increased steady-state cell surface levels of GLUT1 while the L1201A mutant do not (**Figure 5C and 5D**). Finally, we lentivirally expressed dominant negative RAB10(T23N) mutant in parental HeLa cells. This revealed a reduced steady-state surface level of Retromer cargoes GLUT1, CTR1 and KIDINS220 (**Figure 5E and 5F**) (Curnock and Cullen, 2020). Together these data establish the molecular mechanism for the direct association of DENND4A and DENND4C with Retromer and reveal the importance of this interaction for their association to the Retromer retrieval sub-domain. Evidence points to a functional role for these RAB10 GEFs in potentially regulating RAB10-mediated cargo transport through the Retromer sub-domain.

**Figure 5.**
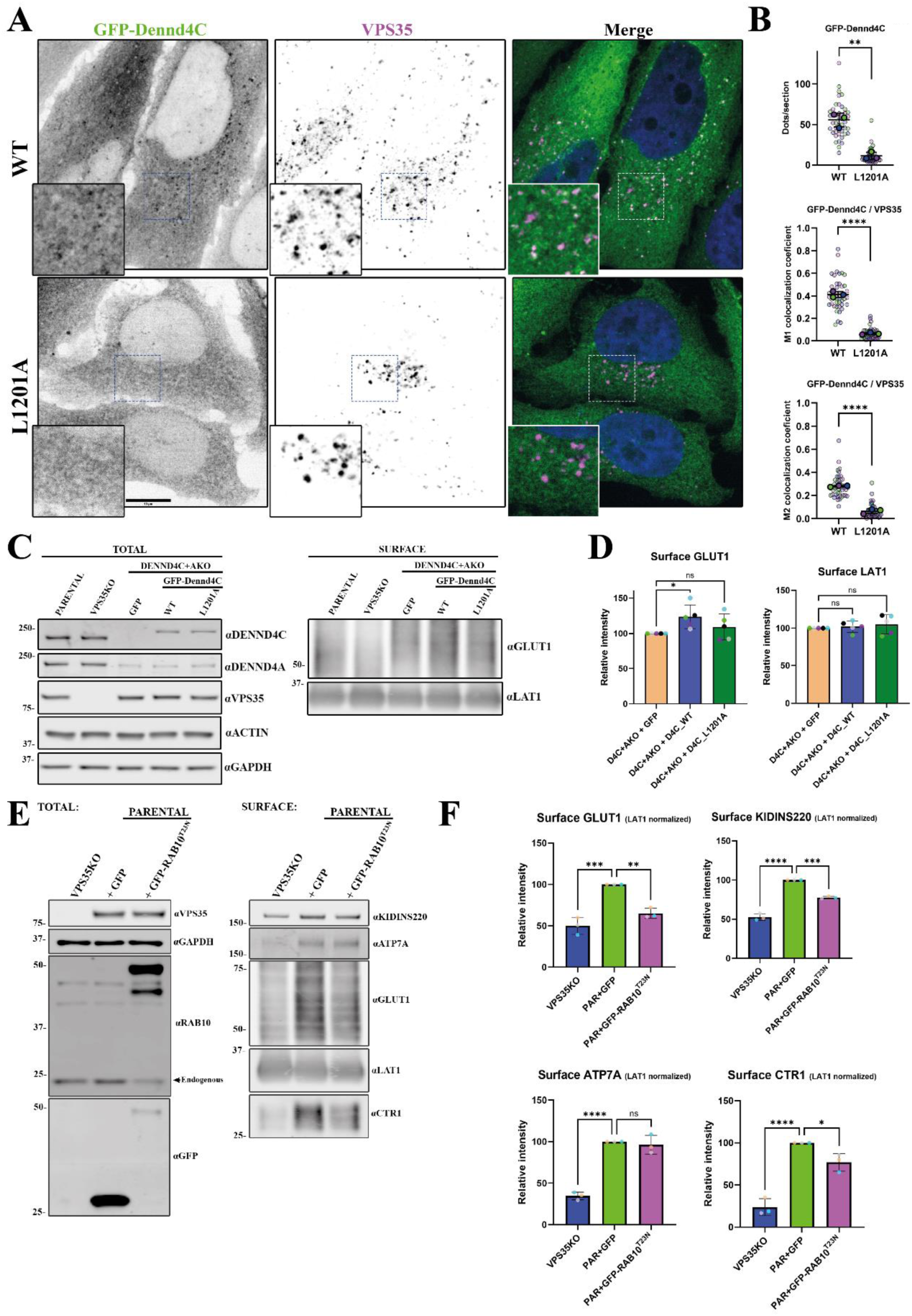
Role of DENND4A/C in Retromer’s cell biology. **(A)** Spinning disk confocal images showing colocalization of stably expressed mouse GFP-Dennd4C with endogenous VPS35 in DENND4C and DENND4A double KO Hela cells; wild-type (WT) and L1201A mutant equivalent to human DENND4C L1204A. Scale bar - 10 μm. **(B)** Quantitation and statistical analysis of number of dots per cell section of GFP-Dennd4C and its colocalization with VPS35 from (A) using ImageJ colocalization JACOP plugin, Manders coefficients M1/M2. n = 3 independent experiments. T-test analysis, data presented as mean values relative to WT and error bars represent SD. **(C)** Analysis of protein surface levels in Hela parental, VPS35KO and DENND4A+C double KO transduced with GFP alone, GFP-Dennd4C WT or L1201A mutant. VPS35KO cell line was used as control for Retromer’s role in GLUT1 recycling. Left panel shows total protein levels while right panel shows cell surface protein fraction after surface biotinylation and streptavidin purification. **(D)** Quantitation and statistical analysis of GLUT1 and LAT1 surface levels in transduced Hela DENND4A+C double KO from experiment (C). n = 5 independent experiments. 1-way ANOVA with Dunnett’s multiple comparison test, data presented as mean values relative to WT and error bars represent SD. **(E)** Analysis of protein surface levels in Hela cells transduced with GFP alone or GFP-RAB10 T23N dominant negative mutant. VPS35KO cell line was used as control for Retromer’s role in protein recycling. Left panel shows total protein levels while right panel shows cell surface protein fraction after surface biotinylation and streptavidin purification. **(F)** Quantitation and statistical analysis of GLUT1, KIDINS220, ATP7A and CTR1 surface levels in Hela cells from experiment (E). Data was normalized to LAT1. n = 3 independent experiments. 1-way ANOVA with Dunnett’s multiple comparison test, data presented as mean values relative to WT and error bars represent SD.

### Interactions of Retromer with the TBC domain family of RAB GAPs

We also identified the RAB10 effector and RAB35 GAP TBC1D13 in both BioID1-VPS35 and VPS35-BioID1 experiments (Davey et al., 2012) (**Figure 1E and 1F**). We therefore modelled the interaction of TBC1D13 with Retromer using AlphaFold2, which consistently predicted a high confidence assembly (**Figure 6A**, ***Supplementary Figure 9***). This revealed a remarkable complex whereby TBC1D13 envelops VPS29 using an extensive clamp structure composed of two loop sequences (residues 80-107 and 164-204). The predicted structure reveals three notable interfaces. The first interface involves loop residues 80-107 containing an HPL motif binding to the canonical surface of VPS29 centred on Leu152 (**Figure 6B view 1**). This is consistent with the loss of TBC1D13 interaction with VPS29 mutants L152E and V154D (**Figure 6C and 6D**). We further validated the importance of this binding site by ITC, with recombinant full-length human TBC1D13 binding to the Retromer complex with an affinity of 6.3 μM, which was largely blocked by the competing cyclic peptide RT-D1 (**Figure 6E and Table 2**). GFP-nanotrap isolation of GFP-tagged TBC1D13 further confirmed binding to Retromer that was lost in the TBC1D13 (P101A) mutant that targeted the HPL motif (**Figure 6F and 6G**). On the opposite face of VPS29 a second interface is formed by residues 164-204 of TBC1D13, which form an extended β-hairpin structure and makes a direct contact with the VPS35 subunit at their extremity (**Figure 6B view 2**). The third notable prediction is that the amino-terminal methionine of VPS29 is largely buried in the structure (**Figure 6B view 3**). This suggests that the full binding affinity for TBC1D13 will require the canonical VPS29 isoform 1, and that the splice variants 2 and 3 with longer amino-terminal sequences could be compromised for TBC1D13 binding. Finally, when expressed in cells full-length GFP-tagged TBC1D13 localised to Retromer decorated endosomes, an endosomal targeting that was completely lost with the TBC1D13(P101A) Retromer-binding mutant (**Figure 6H and 6I**).

**Figure 6.**
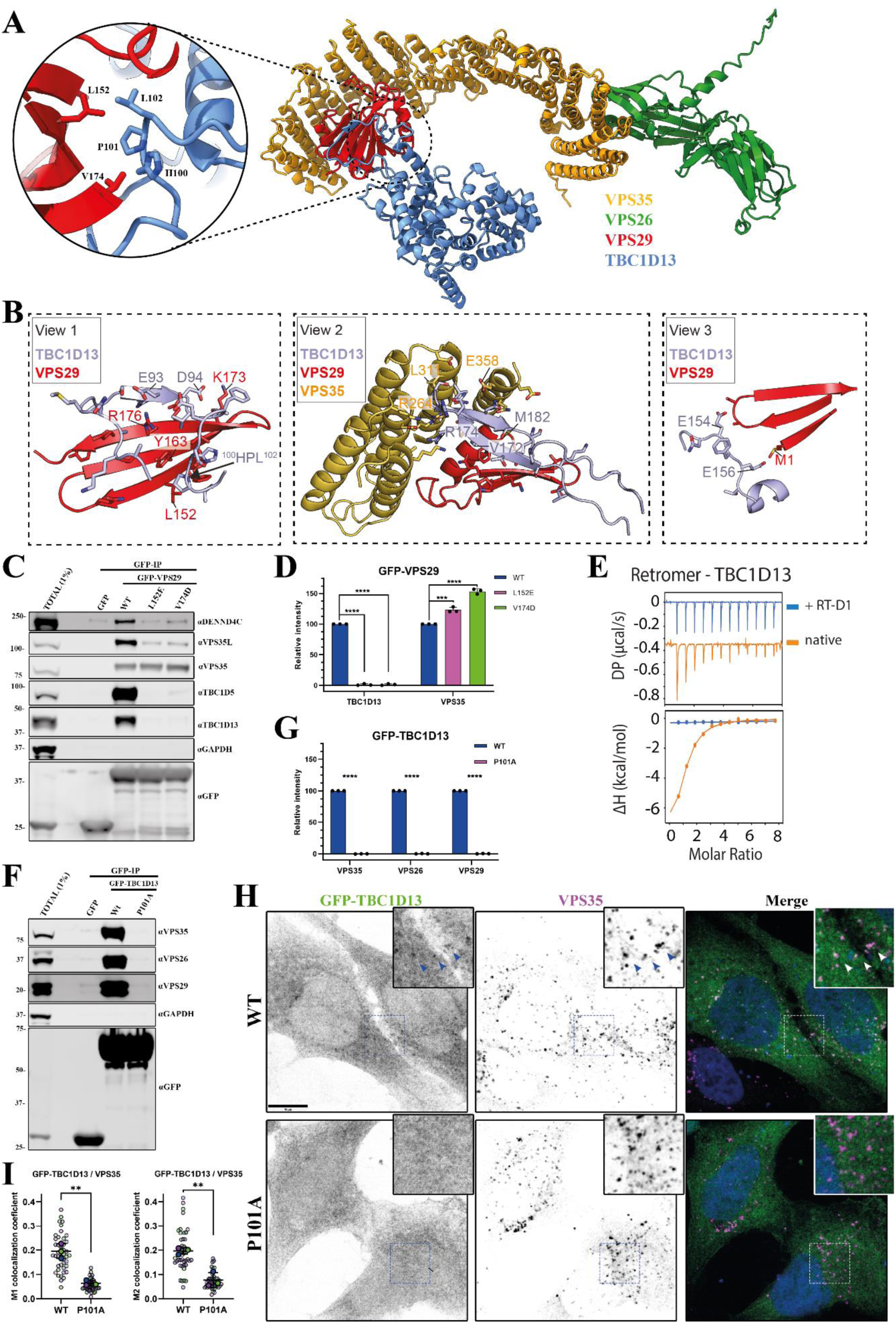
Interactions between TBC1D13 and Retromer complex. **(A)** AlphaFold2 predictive model of TBC1D13 interacting with Retromer (rank 1 prediction). Circular inset shows main binding site consisting of a conserved PL motif (TBC1D13 ^101^PL^102^) interacting with the Leu152 containing hydrophobic cavity of VPS29. **(B)** The TBC1D13 interaction with Retromer involves three notable interfaces. View 1 shows the binding of a PL motif and extended beta-strand to the VPS29 subunit at the canonical site surrounding VPS29 Leu152. View 2 shows an extended beta-hairpin from TBC1D13 on the opposite face of VPS29 that also makes additional contacts with VPS35. Lastly, view 3 shows how VPS29 N-terminal Methionine Met1 (in standard isoform Q9UBQ0) is tightly buried in the complex with TBC1D13. **(C, D)** GFP based co-immunoprecipitation (co-IP) of GFP-VPS29 wild-type (WT) and hydrophobic pocket mutants after transient transfection in HEK293T cells. Quantitation and statistical analysis of relative band intensity for the indicated proteins normalized to GFP band intensity. n = 3 independent experiments. 1-way ANOVA with Dunnett’s multiple comparison test, data presented as mean values relative to WT and error bars represent standard deviation (SD)**. (E)** ITC of TBC1D13 binding to Retromer in the absence or presence of competing cyclic peptide RT-D1. **(F, G)** GFP based co-IP of GFP-TBC1D13 WT and P101A mutant after transient transfection in HEK293T cells. Quantitation and statistical analysis of relative band intensity for the indicated proteins normalized to GFP band intensity. n = 3 independent experiments. T-test analysis, data presented as mean values relative to WT and error bars represent SD. **(H)** Spinning-disk confocal images showing colocalization of transiently expressed GFP-TBC1D13 with endogenous VPS35 in Hela cells; wild-type (WT) and P101A mutant. Cells were fixed ∼15 hrs after plasmid transfection and images were acquired for cells with low to moderate GFP expression. **(I)** Quantitation and statistical analysis of colocalization between GFP-TBC1D13 and VPS35 after manual exclusion of Golgi area and using ImageJ colocalization JACOP plugin, Manders coefficients M1/M2. n = 3 independent experiments. T-test analysis, data presented as mean values relative to WT and error bars represent SD. Scale bar - 10 μm.

**Table 2.**
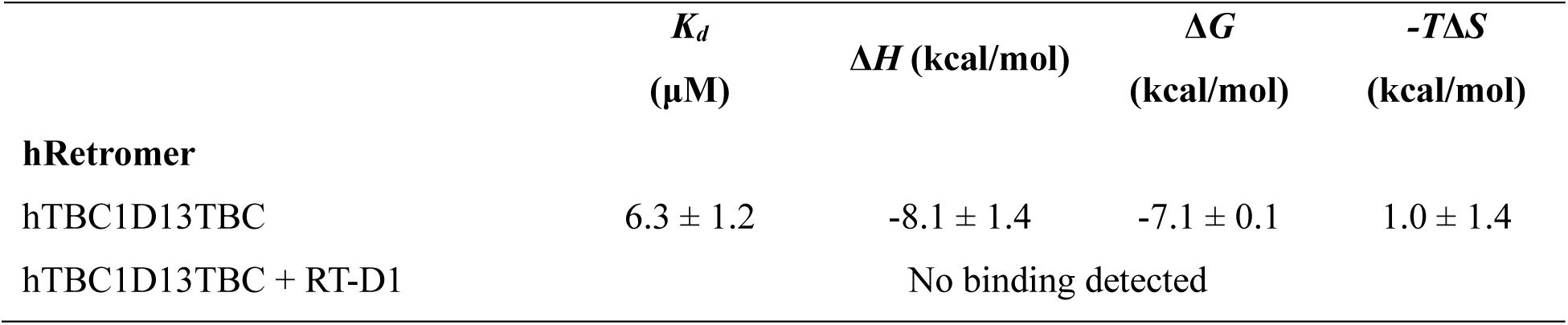
Thermodynamic parameters for the binding of Retromer and TBC1D13 by ITC.

| | $K_d$<br>(μM) | $\Delta H$ (kcal/mol) | $\Delta G$<br>(kcal/mol) | $-T\Delta S$<br>(kcal/mol) |
| --- | --- | --- | --- | --- |
| **hRetromer** |  |  |  |  |
| hTBC1D13TBC | 6.3 ± 1.2 | -8.1 ± 1.4 | -7.1 ± 0.1 | 1.0 ± 1.4 |
| hTBC1D13TBC + RT-D1 |  | No binding detected |  |  |

### In silico screening identifies additional RAB GEFs and RAB GAPs as Retromer accessory proteins

To further explore the ability of Retromer to potentially associate with RAB GEFs and RAB GAPs we performed an *in-silico* screen of all human DENND and TBC domain-containing family members. We applied AlphaFold2 and Alphafold3 to screen Retromer using an approach similar to our previous studies of the Mint interactome (**Figure 7A and 7B**, ***Supplementary Figure 10***) (Weeratunga et al., 2023). To carry out confident prediction over non-specific associations, Alphafold modelling was carried out using Retromer, a Retromer sub-complex or individual Retromer subunits against TBCs/DENNDs (**Figure 7A and 7B**). The IPTM score generated by different runs were combined to pick up those that showed consistent binding to specific Retromer subunits with high IPTM score over non-specific interactors. Using this approach, among the 14 human DENND family members only DENND4A and DENND4C were predicted to bind VPS35:VPS29 with high confidence (**Figure 7A and 7B**). Further validation using pDOCKQ and SPOC, confirmed that both DENND4A and DENNDC associated with the VPS29 subunit (**Figure 7B)**. Interestingly, the poorly characterized DENND11 also possessed a PL motif but did not display sufficient confidence binding score with Retromer subunits and expressed GFP-DENND11 showed weak association with Retromer when immuno-isolated from HEK cells, independently of the PL motif (***Supplementary Figure 10C***).

**Figure 7.**
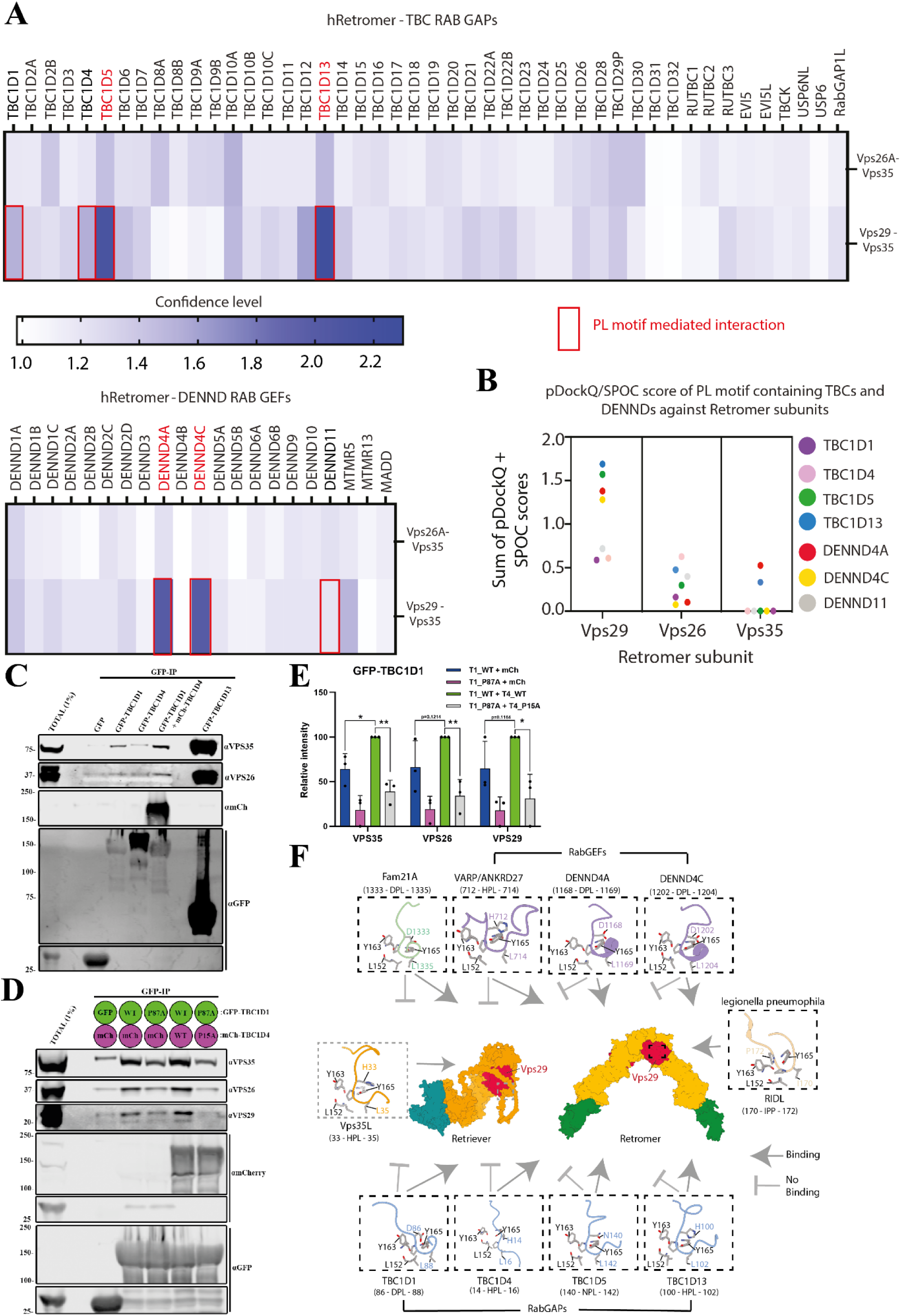
Retromer as a hub for RAB GTPase switch regulation: VPS29 interaction screen and TBC1D1/4 interaction with Retromer. **(A, B)** Retromer interactions with TBC GAP and DENND GEF proteins across the human proteome. VPS26A-VPS35 and VPS29-VPS35 assemblies or individual VPS29, VPS26A and VPS35 proteins were screened against all human proteins with TBC and DENN domains using AlphaFold2. The confidence of potential interactors was scored based on the sum of the interfacial PTM score (iPTM) averaged from three models, pDOCKQ and SPOC scores. In TBC domain GAPs, TBC1D5 and TBC1D13 shows high confidence. In DENN domain GEFs, DENND4A and DENND4C were confidently predicted to bind to VPS29/Retromer. Others found to possess a PL motif but show low confidence in association with VPS29 binding are TBC1D1, TBC1D4 and DENND11. **(C)** GFP based co-immunoprecipitation (co-IP) of GFP-TBC1D1, GFP-TBC1D4 or GFP-TBC1D13 after transient transfection in HEK293T cells. Co-expression of GFP-TBC1D1 with mCherry-TBC1D4 increased pull-down of Retromer complex subunits. **(D, E)** GFP based co-IP after transient transfection in HEK293T cells of wild-type (WT) and PL interacting-motive mutant versions of GFP-TBC1D1 (P87A) and mCherry-TBC1D4 (P15A). Quantitation and statistical analysis of relative band intensity for the indicated proteins normalized to GFP band intensity. n = 3 independent experiments. T-test analysis, data presented as mean values relative to WT and error bars represent SD. **(F)** Schematic summarizing the interactions of Retromer with the identified TBC GAPs and DENND GEFs and the similarity with binding to the FAM21 subunit of the WASH complex and the RidL protein from legionella. The intramolecular occlusion of the equivalent binding site in VPS29 when assembled in the Retriever complex prevents Retriever from binding to any of these proteins through these specific mechanisms.

As for TBC domain-containing GAPs, confident prediction was observed for VPS29 with TBC1D13 and the previously characterised TBC1D5 (Jia et al., 2016). None of the TBCs and DENNDs showed high binding confidence to the VPS26A:VPS35 interface (**Figure 7A**). Screening among the TBCs with low confidence in binding to Retromer, VPS29 was predicted to bind PL motif in both TBC1D1 and TBC1D4, GAPs for RABs that include RAB10 (Mîinea et al., 2005) (**Figure 7B**). Both TBCs reveal binding to the conserved hydrophobic cavity of VPS29 through their PL motif (^86^DPL^88^ in TBC1D1 and ^14^HPL^16^ in TBC1D4) but with limited interacting residues (***Supplementary Figure 11***). Indeed, when expressed as GFP fusions and GFP-trap immuno-isolated GFP-TBC1D1 and GFP-TBC1D4 showed limited binding to endogenous Retromer (**Figure 7C**). Interestingly, further Alphafold2 screening in combination with SPOC and pDOCKQ of the PL motifs containing TBCs/DENNDs, revealed that TBC1D1 and TBC1D4 are likely to form a heterodimer through their C-terminal tails (***Supplementary Figure 12*)**. Similar dimerization also observed in TBC1D5, showing a small region within the C-terminal disordered tail likely involved in homodimerization (***Supplementary Figure 12*)**. In TBC1D1 and TBC1D4, the disordered tail region after the TBC domain likely responsible for both homo- and heterodimerization (***Supplementary Figure 12*)**. To examine whether heterodimerization enhances binding to Retromer, we co-expressed GFP-TBC1D1 with mCherry-TBC1D4 and performed GFP-nanotrap immuno-isolation. This established that these RAB GAPs do indeed form heterodimers and that this appears to enhance association with endogenous Retromer (**Figure 7D and 7E**). Under these conditions mutagenesis of both PL motifs, TBC1D1(P87A) and TBC1D4(P15A) significantly reduce Retromer association (**Figures 7D and 7E**).

Overall, with multiple copies of Retromer being enriched within the retrieval sub-domain and lower concentrations across the limiting endosomal membrane, these data provide mechanistic evidence for a model where Retromer associates with selected RAB GEFs and RAB GAPs to regulate the activity of key endosomal RAB identity cues (**Figure 7F**). These include RAB7, RAB10, RAB21, and RAB35 and most likely other RAB GTPases given the broad spectrum of RAB substrates for some of these regulators.

## DISCUSSION

Central to achieving a thorough understanding of the mechanistic basis and functional significance of endosomal cargo retrieval and recycling will be a detailed appreciation of the components of the retrieval sub-domain, their organisation, and their dynamic regulation (Cullen and Steinberg, 2018; Norris and Grant, 2020). Here, we have engineered proximity proteomic technology to provide a quantitative approach for the biochemical identification of the components of the endosomal Retromer and Retriever demarcated retrieval sub-domain. This has revealed quantitative *in-cellulo* biochemical data supportive of the structure and organisation of Retromer and Retriever and a wealth of new molecular insight to pave the way to a greater mechanistic understanding of Retromer and, more generally, retrieval sub-domain structure, organisation and function in human health and disease.

Through designing our BioID strategy into a single proximity proteomic analysis we have directly compared the quantitative proximity signatures of the HRS-demarcated degradative sub-domain with the Retromer and Retriever defined retrieval sub-domain. This has established a clear biochemical separation in the proximity profile of these degradative and retrieval sub-domains, entirely consistent with imaging data of the spatial separation of these sub-domains on the limiting membrane of individual endosomes (Puthenveedu et al., 2010). When comparing Retromer and Retriever our data suggests that although these heterotrimeric complexes do not appear at steady-state to reside in proximity to one another, they are connected through shared proximity profiles with the WASH complex. This is entirely consistent with the known direct association of Retromer with the FAM21 subunit of the WASH complex (Harbour et al., 2012; Jia et al., 2012; Guo et al., 2024; Romano-Moreno et al., 2024), and the less well-described coupling of Retriever to the WASH complex by means of the CCC complex (Bartuzi et al., 2016). It is also consistent with the essential role of the WASH complex and its ability to regulate localised branched F-actin dynamics in the organisation of the Retromer and Retriever demarcated retrieval sub-domain (Puthenveedu et al., 2010; Harbour et al., 2012; Jia et al., 2012; Gomez et al., 2012; Bartuzi et al., 2016; Lee et al., 2016; Simonetti and Cullen, 2019).

By strategic engineering of the BioID1 enzyme to the amino-terminus and carboxy-terminus of the extended core VPS35 α-solenoid of Retromer we have provided the first *in-cellulo* quantitative biochemical data supporting the arch-like organisation of the membrane associated Retromer dimers as observed by cryo-ET of the *in vitro* reconstituted liposome associated Retromer (Kovtun et al., 2018; Leneva et al., 2021). The proximity profile of the amino-terminal BioID1 tagged VPS35 is consistent with the juxtaposed position of this region to the cytoplasmic facing endosomal membrane, while the carboxy-terminal BioID1 proximity proteome reflects the membrane distal localisation at the apex of Retromer arches. These data therefore establish biochemical signatures for the inner layer (*e.g.* RAB7, SNX3, and various cargo proteins) and outer layer (*e.g.* TBC1D5, ANKRD27, DNAJC13 and the WASH complex) of proteins associated with the Retromer coat complex and/or transiting through the Retromer retrieval sub-domain. Finally, for the corresponding core VPS35L α-solenoid of Retriever such a clear separation between the amino-terminal and carboxy-terminal proximity proteomes is not observed. The recently described structure of Retriever and its association with the CCC complex to form the Commander super-assembly provides the molecular explanation for these data (Healy et al., 2023; Boesch et al., 2024; Laulumaa et al., 2024). Here, the amino-terminus and carboxy-terminus of VPS35L reside in proximity to one another. Our proximity proteomic data therefore provides quantitative *in-cellulo* evidence consistent with the organisational features of Retriever and its assembly into Commander.

In addition to the validation of established structural and organisational features of Retromer and Retriever and their known accessory proteins, our data-rich proximity proteomic resource provides a wealth of new molecular details of endosomal cargo retrieval and recycling. Our identification and preliminary validation of the augmin complex within the Retromer proximity proteome suggests a potential localised regulation of microtubule assembly during endosomal function or during a specialised role such as in cell division (Akhmanova and Kapitein, 2022). Retromer has previously been linked to the γTuRC through its accessory protein ANKRD50 (Kvainickas et al., 2017), and endosomal dynamics and endosomal cargo retrieval and recycling are heavily reliant on plus-end and minus-end directed microtubule motor proteins (Wassmer et al., 2009; Hunt et al., 2013; Jongsma et al., 2023). Exploring a potential link of the endosomal Retromer retrieval sub-domain with the regulation of localised microtubule dynamics, especially within non-mitotic neurons, certainly warrants further investigation.

The Retromer proximity proteome has highlighted a close association with regulators of small GTPases including well-established Retromer accessory proteins TBC1D5 and ANKRD27 (Jia et al., 2016; Crawley-Snowdon et al., 2020). We have now extended these regulators to include DENND4A and DENND4C, TBC1D13, and a TBC1D1/TBC1D4 heterodimer, and in so doing established new mechanistic links between Retromer and the switching of RAB10 and RAB35; two RAB GTPases previously linked to the regulation of endosomal cargo recycling (Chen et al., 2006; Glodowski et al., 2007; Sato et al., 2008; Rahajeng et al., 2012; Wang et al., 2016; Klinkert and Echard, 2016; Liu et al., 2018; Etoh and Fukuda et al., 2019; Khan et al., 2022). Our structural dissection of Retromer’s association to these RAB10 and RAB35 regulators has provided the mechanism to link sequence-dependent cargo selection and retrieval sub-domain organisation with the regulation of these switches during endosomal cargo retrieval and recycling. In all cases, the accessory protein directly associates with Retromer through presentation of PL motifs to the same hydrophobic pocket in VPS29 that accommodates PL motifs from TBC1D5, ANKRD27 and the FAM21 subunit of the WASH complex (Jia et al., 2016; Crawley-Snowdon et al., 2020; Guo et al., 2024; Romano-Moreno et al., 2024). Secondary associations with other features of VPS29 or VPS35 are required to stabilise the overall association to Retromer. With evidence that Retromer is required for the endosomal targeting of these RAB regulators, these data reveal a far greater complexity in the role of Retromer in regulating selected RAB GTPase than previously appreciated. In addition to its scaffolding roles in associating with cargo adaptors for the sequence-dependent recognition of cargo proteins and the WASH complex for aiding retrieval sub-domain organisation, Retromer should therefore also be viewed as a major endosomal hub for controlling the activation status and switching of specific RAB GTPases. This feature of Retromer appears to be unique in that, because of its structural architecture (Healy et al., 2023; Boesch et al., 2024; Laulumaa et al., 2024), Retriever is unable to associate with any of these RAB regulators.

Consistent with the direct association of Retromer to the RAB10 GEFs DENND4A and DENND4C and the RAB GAPs TBC1D1 and TBC1D4, functional analysis has identified a role for RAB10 in the endosomal retrieval and cell surface recycling of a classic Retromer cargo, the glucose transport GLUT1. It is important to note however, that while this cargo transport phenotype is quantifiable and statistically significant the level of phenotypic penetrance is far less than observed upon perturbation of SNX27 (the cargo adaptor for GLUT1) or Retromer (Steinberg et al., 2013). This may reflect a level of redundancy in the routes that GLUT1 may take to recycle back to the cell surface once the essential SNX27 and Retromer orchestrated retrieval fate decision has been made. Indeed, an element of this redundancy may stem from Retromer’s hub-like ability to regulate several RABs linked to endosomal cargo recycling.

Finally, it is worth noting that several RABs identified as being controlled by the Retromer hub are substrates for LRKK2 mediated phosphorylation in response to disrupted lysosomal homeostasis (Ito et al., 2016; Steger et al., 2016; Taylor and Alessi, 2020). As the Parkinson’s disease associated VPS35(D620N) mutation leads to enhanced LRRK2 activation and target RAB phosphorylation (Mir et al., 2018), it is tempting to speculate that Retromer’s role as a RAB regulatory hub may constitute a feedback controller in integrating lysosomal homeostasis with endosomal retrieval sub-domain function. Exploring this potential point of connectivity may provide new insight into Retromer’s neuroprotective role in Alzheimer’s disease, Parkinson’s disease and other neurodegenerative conditions.

### Limitations of the study

While we have utilised CRISPR-Cas9 knockout and titrated lentiviral transduction of BioID tagged rescue transgenes to allow as near to endogenous expression as technically possible, BioID tagging of the endogenous gene loci will allow endogenous tagging of proteins to further validate and extend the present findings. The relatively poor kinetics of proximity labelling by BioID1 has limited our study to a steady-state analysis. While this has established the correct targeting strategy for each chosen protein and has revealed new molecular insight, replacement of the BioID1 enzyme with more kinetically rapid proximity labelling reagents will allow for time-resolve proximity proteomics. This will provide temporal access to the dynamics of the retrieval sub-domain make-up, organisation, and adaptation during stimulated cargo sorting. Finally, while our study has identified new links between the retrieval sub-domain and SNARE proteins, the augmin complex, caveolae, and additional functionally interesting proteins, a great deal of further experimentation will be required to validate their localisation and functional role within the retrieval sub-domain.

## MATERIALS AND METHODS

### Antibodies

Primary antibodies: β-Actin (Sigma-Aldrich, A1978; 1:5000 WB, 1:1000 IF), GAPDH (Sigma-Aldrich, G9545; 1:5000 WB), GAPDH (Sigma-Aldrich, G8795; 1:5000 WB), EEA1 (Cell Signalling; 3288S; 1:200 IF), GFP (Roche; 11814460001; clones 7.1/13.1; 1:1000 WB), mCherry/RFP (Abcam, 167453; 1:1000 WB), FLAG-M2 (Sigma, F1804; 1:1000 WB, 1:200 IF), GLUT1 (Abcam; ab115730; 1:1000 WB; 1:400 IF), LAMP1 (Developmental Studies Hybridoma Bank; AB_2296838; clone H4A3; 1:400 IF), VPS29 (Santa Cruz; D-1; sc-398874; 1:500 WB), VPS35 (Abcam, ab97545; 1:1000 WB, 1:400 IF), VPS35 (Antibodies.com, A83699; 1:400 IF), VPS26 (Abcam, ab23892; 1:1000 WB), VPS35L (Abcam, ab97889; 1:1000 WB), VPS26C (Sigma-Aldrich, ABN87; 1:1000 WB), VPS35L (Invitrogen, PA5-28553; 1:200 IF), SNX1 (BD Transduction Lab, 611482; 1:1000 WB, 1:200 IF), SNX1 (Proteintech, 10304-1-AP; 1:200 IF), SNX2 (BD Transduction Lab, 611308; 1:1000 WB), SNX5 (Abcam, ab180520; 1:1000 WB), SNX6 (Santa Cruz, sc-365965; 1:1000 WB), SNX17 (Proteintech, 10275-1-AP; 1:1000 WB), SNX17 (Sigma, HPA043867, 1:100 IF), SNX27 (Proteintech, 16329-1-AP; 1:1000 WB), SNX27 (Abcam, ab77799; 1:100 IF), HRS/HGS (Enzo Life Sciences, ALX-804-382-C050; 1:2000 WB, 1:100 IF), STAM (Proteintech, 12434-1-AP; 1:2000 WB, 1:100 IF), FAM21 (Gift from Dan Billadeau; 1:2000 WB), Integrin-α5 (Abcam, ab150361; 1:1000 WB, 1;200 IF), DENND4C (Sigma-Aldrich, HPA014917, 1:1000 WB, 1:500 IF), DENND4C (StressMarq Bioscience, SMC-610, 1:100 IF), DENND4A (Abcam, ab117758, 1:500 WB), HAUS2 (ThermoFisher, PA5-31258; 1:500 WB), HAUS6 (Proteintech, 16933-1-AP; 1:1000 WB), HAUS8 (Abcam, ab95970; 1:250 WB), CAV1 (Proteintech, 16447-1-AP; 1:1000 WB), CAVIN1 (Proteintech, 18892-1-AP; 1:1000 WB), TBC1D13 (ThermoFisher, PA5-61110; 1:2000 WB, 1:200 IF), ARHGAP1 (Proteintech, 11169-1-AP; 1:1000 WB, 1:100 IF), VAMP3 (Proteintech, 10702-1-AP; 1:1000 WB, 1:200 IF), NF1 (Proteintech, 27249-1-AP, 1:1000 WB), ARFGEF1 (Abcam, A44423,1:1000 WB), TBC1D4 (Proteintech, 68063), KIDINS220 (Proteintech, 21856-1-AP, 1:1000 WB), ATP7A (Santa Cruz, sc-376467, 1:1000 WB), CTR1 (Abcam, ab129067, 1:1000 WB), LAT1 (Cell Signaling, 5347s, 1:1000 WB), TBC1D5 (Abcam, 203896, 1:1000 WB), VARP/ANKRD27 (Proteintech, 24034-1-AP, 1:1000 WB), RAB10 (Abcam, ab237703, 1:1000 WB, 1:200 IF). Secondary antibodies for WB: 680nm and 800nm donkey anti-mouse and anti-rabbit fluorescent secondary antibodies (Invitrogen, A-21057, A3275; 1:20,000). Secondary antibodies for IF: 405nm, 488nm, 568nm and 647nm AlexaFluor-labelled anti-mouse, anti-rabbit or anti-goat (Invitrogen; 1:400).

### Plasmids and molecular cloning

cDNA codifying for the proteins expressed were cloned into pEGFP/mCherry-C1/N1 transient vectors or pLVX lentiviral vectors by restriction enzyme digestion and ligation or Gibson Assembly reaction. Sequence for BioID1 preceded by FLAG tag was fused by a flexible linker to the N or C terminus of selected proteins and incorporated in pLVX-CMV-MCS-PGK-Puro vector. CRISPR-Cas9 gRNAs were designed as a couple of opposing DNA primers which were phosphorylated, annealed and cloned into pSpCas9(BB)-2A-Puro (pX459) BbsI digested. Resulting plasmids were transformed in XL1-Blue (Agilent) or NEB 5-alpha HE (NEB) *E. coli* strains and grown on selecting media, purified and validated by sanger sequencing. gRNAs sequences are shown in Table S1 and list of plasmids in Table S2.

For bacterial expression, full-length human VPS35 and VPS26A with an N-terminal Hi-tag were cloned into pET28a vector as described previously (Norwood et al., 2011; Chen et al., 2021). Human VPS29 was cloned into pGEX4T-2 vector with a linker between the start of the gene and the thrombin recognition site to facilitate the tag cleavage. DNA encoding the bacteria expression optimized human full-length TBC1D13 was synthesized and cloned into pGEX6P-1 vector by Gene Universal. All DNA constructs were verified using DNA sequencing.

### Cell culture, transfection and lentiviral production

HeLa and HEK293T cell lines were sourced from ATCC. Authentication was from the ATCC and additionally, HeLa line was authenticated by Eurofins services. Cells were grown in DMEM (Sigma-Aldrich) supplemented with 10% (v/v) FCS (Sigma-Aldrich) and penicillin/streptomycin (Gibco, USA) and grown in humidified incubators at 37°C, 5 % CO_2_. FuGENE HD (Promega, USA) or Lipofectamine LTX (Invitrogen) were used for transient transfection of DNA according to the manufacturer’s instructions. Alternatively, for immunoprecipitation or viral production, cells were transfected with polyethylenimine (PEI) (Sigma-Aldrich) in 3:1 PEI:DNA weight ratio.

For the generation of CRISPR-Cas9 KO HeLa cells, cells were transfected with pX459 plasmid coding for the gRNA against the gene of interest as well the puromycin resistance. One day after transfection, cells were subjected to puromycin selection for 24 hours. After 3-4 days recovering without puromycin, the pooled population of cells was subjected to lysis and WB to confirm the KO. Once the gRNA was validated, the experiment was repeated, but after puromycin selection, cells were diluted for clonal selection in 96-well plates with IMDM (Sigma-Aldrich). After 3-4 weeks clonal populations were escalated and checked by WB. VPS35KO, SNX27KO, SNX17KO and VPS35LKO clonal cell lines used in this study were previously characterized (Simonetti et al., 2022; McNally et al., 2017; Healy et al., 2023).

For transient GFP/mCherry-based immunoprecipitations, HEK293T cells in 15 cm plates at 70% confluency were transfected with GFP/mCherry plasmids using PEI procedure, and after 24-48 hrs, cultures were lysed for protein purification. For lentiviral production, HEK293T cells in 15cm plates at 70% confluency were co-transfected with the pLVX and the helper plasmids PAX2 and pMDG2 using PEI. After 4 hrs incubation the media was replaced and 48hs after transfection the media containing the virus was harvested, spun down and filtered through 0.45 µm filters before being aliquoted and stored at −80°C.

### Immunoprecipitation and quantitative western blot analysis

For every kind of protein extraction, buffers were pre-chilled, and cells and samples placed on ice. For basic western blot (WB), cells were lysed in ∼ pH 7.2 phosphate buffered saline (PBS) with 1% (v/v) Triton X-100 and protease inhibitors (Thermo Fisher, A32955). The protein concentration was determined with a bicinchoninic acid (BCA) assay kit (Thermo Fisher, 23225) and equal amounts were denaturised (LDS Sample buffer (Thermo Fisher, NP0008), 3% β-mercaptoethanol (Sigma-Aldrich, M6250), 10 min at 95°C) and resolved by SDS-PAGE on NuPAGE 4-12% precast gels (Invitrogen, NP0336BOX). Protein samples were then transferred onto methanol-activated polyvinylidene fluoride (PVDF) membrane (Immobilon-FL membrane, pore size 0.45μm; Millipore, IPFL00010). Membrane was blocked, then sequentially labelled with primary and secondary antibodies. Fluorescence detected by scanning with a LI-COR Odyssey scanner and Image Studio analysis software (LI-COR Biosciences). Typically, we performed WB in where a single blot is simultaneously probed with antibodies against 2 proteins of interest (distinct antibody species) followed by visualization with the corresponding secondary antibodies conjugated to distinct spectral dyes.

For GFP/mCherry-based co-immunoprecipitations (co-IP), cells were lysed in IP buffer (50 mM Tris-HCl, 0.5% (v/v) NP-40, and protease inhibitors) and centrifuged 10 min at 16.000xg. An aliquot of the cleared lysate was retained to represent the whole cell fraction and the rest added to pre-washed (in IP buffer) GFP/mCherry-trap beads (ChromoTek, gta-20/rta-20) for rocking 1 hr at 4°C. After, beads were washed 3 times with IP buffer by rounds of re-suspension and pelleting. Finally, buffer was removed, and beads were then either stored at −20°C or processed for SDS-PAGE analysis. For proteomic analysis, detergent in final washes was sequentially removed, and samples processed immediately.

### BioID1 protein-proximity labelling

Cells stably expressing the protein of interest tagged with BioID1 were grown in 15 cm plates. When confluency reached 70%, media was changed by DMEM supplemented with 50 μM biotin (Sigma-Aldrich) and incubated typically for 24 hrs. Cells were washed twice in PBS and lysed in RIPA buffer (50 mM Tris-HCl pH 7.5, 150 mM NaCl, 0.1% SDS (v/v), 0.5% (w/v) Sodium Deoxycholate, 1% (v/v) Triton X-100 and protease inhibitors (Roche, 05892970001)). Lysates were centrifuged 10 min at 16.000xg and the supernatants were collected. The protein concentration was determined with a BCA assay kit. An aliquot of the cleared lysate was retained to represent the whole cell fraction and the rest added to pre-washed (in RIPA buffer) streptavidin sepharose beads (Sigma-Aldrich, Cytiva 17511301) for rocking 2 hrs at 4°C. Later, beads were sequentially washed by re-suspension and pelleting: twice in RIPA buffer, once in 1 M KCl, once in 100 mM Na_2_CO_3_, once in 2 M Urea in 10 mM Tris-HCl pH 8 and twice in RIPA buffer. Finally, whole cell fractions and beads were either stored at −20°C or denaturised (LDS Sample buffer, 3% β-mercaptoethanol, 20 mM DTT (Thermo Fisher, 20290), 2 mM biotin, 10 min at 95°C) and resolved by SDS-PAGE. For proteomic analysis, beads were washed once more with RIPA buffer without detergents, and samples processed immediately.

### Surface biotinylation

All steps were carried out on ice to prevent surface protein internalisation. Just prior to starting the biotinylation labelling, membrane impermeable Sulfo-NHS-SS-Biotin (Thermo Fisher, 21331) was dissolved at a final concentration of 0.2 mg/ml in PBS adjusted to pH 7.8. Cells were washed twice in cold PBS to stop endocytosis before being incubated with Sulfo-NHS-SS-Biotin for 20 min. Later, cells were washed twice in quenching buffer (50 mM Tris-HCl pH 7.5, 100 mM NaCl) and incubated on it for 10 min. Then, cells were lysed in PBS, 1% Triton X-100 plus protease inhibitor cocktail (Thermo Fisher, A32955). Lysates were centrifuged 10 min at 16.000xg and the supernatants were collected. The protein concentration was determined by BCA. An aliquot of the cleared lysate was retained to represent the whole cell fraction and the rest added to pre-washed (in PBS plus 1% Triton) streptavidin sepharose beads for rocking 1 hr at 4°C. Afterwards, beads were sequentially washed by re-suspension and pelleting: three times in PBS plus 1% Triton plus 1.2 M NaCl. Finally, whole cell fractions and beads were either stored at −20°C or denaturised (LDS Sample buffer, 3% β-mercaptoethanol, 20 mM DTT, 2 mM biotin, 10 min at 95°C) and resolved by SDS-PAGE.

### Immunofluorescence Microscopy and Analysis

HeLa cells were seeded onto 13 mm coverslips. Cells were fixed in 4% (v/v) paraformaldehyde (PFA) (Pierce, 28906) in PBS for 20 min and permeabilised in 0.1% (v/v) Triton X-100 in PBS (Sigma-Aldrich), or in 0.1% (w/v) saponin (sigma-Aldrich) in PBS when labelling LAMP1 compartments, for 5 minutes followed by blocking with 2% (w/v) BSA (Sigma, 05482) in PBS for 30 min. Coverslips were stained with primary antibodies for 1 hr followed by secondary antibodies for 30 minutes, then mounted onto glass microscope slides with Fluoromount-G (Invitrogen, 00-4958-02).

Confocal microscope images were taken on a Leica SP5-II confocal laser scanning microscope attached to a Leica DMI 6000 inverted epifluorescence microscope or a Leica SP8 confocal laser scanning microscope attached to a Leica DM l8 inverted epifluorescence microscope (Leica Microsystems), with a 63x UV oil immersion lens, numerical aperture 1.4 (Leica Microsystems, 506192). For the Leica SP8 microscope, ‘lightning’ adaptive image restoration was used to generate deconvolved representative images. Colocalization analysis was performed using Volocity 6.3 software (PerkinElmer) or JACOP plugin from ImageJ-FIJI software. Spinning disk high-resolution images were acquired on an IXplore SpinSR system (Olympus) consisting of an IX83 microscope frame with a CO_2_ and temperature chamber (Olympus), SoRa W1 twin cam spinning disk unit (Yokogawa) and two back thinned Fusion BT sCMOS cameras (Hamamatsu). A 488nm or 561nm laser provided excitation for the green and red channel respectively. The SoRa disk with additional 3.2x magnification changer was used for imaging with excitation light focused to the sample using a 60x/1.4NA oil immersion lens. The system operated in simultaneous twin cam mode and the desired fluorescence was selected through use of either a 525/50nm or 617/73nm bandpass filter for the green or red channel respectively.

### TMT Labelling and High pH reversed-phase chromatography

Proteomic experiments were performed with isobaric tandem mass tagging (TMT) coupled to nanoscale liquid chromatography joined to quantitative tandem mass spectrometry (nano-LC-MS/MS). Bead samples were immediately reduced with 10 mM TCEP (55°C for 1 h), alkylated with 18.75 mM iodoacetamide (room temperature for 30 min) and then digested from the beads with trypsin (2.5 μg trypsin; 37°C, overnight). After the digestion, the resulting peptides were labelled with TMT 10 or 8 plex reagents according to the manufacturer’s protocol (Thermo Fisher). The samples were pooled and evaporated to dryness, resuspended in 5% formic acid and next desalted using a SepPak cartridge according to the manufacturer’s instructions (Waters^TM^). Eluate from the SepPak cartridge was again evaporated to dryness and resuspended in buffer A (20 mM NH₄OH, pH 10) before to be fractionated by high pH reversed-phase chromatography using an Ultimate 3000 liquid chromatography system (Thermo Fisher). Shortly, the sample was loaded onto an XBridge BEH C18 Column (130 Å, 3.5 μm, 2.1 mm X 150 mm, Waters^TM^) in buffer A and the peptides were eluted with an increasing gradient of buffer B (20 mM Ammonium Hydroxide in acetonitrile, pH 10) from 0-95% over 60 min. 5 fractions were generated, evaporated to dryness and finally resuspended in 1% formic acid to get them ready for analysis by nano-LC MSMS using an Orbitrap Fusion Tribrid mass spectrometer (Thermo Fisher).

### Nano-LC Mass Spectrometry

The resulting fractions were further fractionated with an Ultimate 3000 nano-LC system in line with an Orbitrap Fusion Tribrid mass spectrometer (Thermo Scientific). Concisely, peptides in 1% (vol/vol) formic acid were injected onto an Acclaim PepMap C18 nano-trap column (Thermo Scientific). After washing with 0.5% (v/v) acetonitrile and 0.1% (v/v) formic acid, peptides were resolved on a 250 mm × 75 μm Acclaim PepMap C18 reverse phase analytical column (Thermo Scientific) over a 150 min organic gradient, including 7 gradient segments (1-6% solvent B over 1 min, 6-15% B over 58 min, 15-32% B over 58 min, 32-40% B over 5 min, 40-90%B over 1 min, held at 90% B for 6 min and then reduced to 1% B over 1 min) at a flow rate of 300 n/min. Solvent A was 0.1% formic acid and Solvent B was aqueous 80% acetonitrile in 0.1% formic acid (v/v). Peptides were ionized by nano-electrospray ionization at 2.0kV using a stainless-steel emitter with an internal diameter of 30 μm (Thermo Scientific) and a capillary temperature of 275°C. All spectra were acquired using an Orbitrap Fusion Tribrid mass spectrometer controlled by Xcalibur 3.0 software (Thermo Scientific) and operated in data dependent acquisition mode using an SPS-MS3 workflow. FTMS1 spectra were collected at a resolution of 120,000, with an automatic gain control (AGC) target of 200,000 and a maximum injection time of 50 ms. Precursors were filtered with an intensity threshold of 5,000, according to charge state (to include charge states 2-7) and with monoisotopic peak determination set to peptide. Formerly interrogated precursors were omitted using a dynamic window (60 s +/- 10 ppm). The MS2 precursors were isolated with a quadrupole isolation window of 1.2 m/z. ITMS2 spectra were collected with an AGC target of 10,000, CID collision energy of 35% and max injection time of 70 ms. For FTMS3 analysis, the Orbitrap was operated at 50,000 resolution with an AGC target of 50,000 and a maximum injection time of 105 ms. Precursors were fragmented by high energy collision dissociation (HCD) at a normalised collision energy of 60% to ensure maximal TMT reporter ion yield. Synchronous Precursor Selection (SPS) was enabled to include up to 10 MS2 fragment ions in the FTMS3 scan.

### Statistics and bioinformatic analysis

Proteomic raw data files were processed and quantified using Proteome Discoverer software v2.1 (Thermo Scientific) with the SEQUEST HT algorithm and searching against the UniProt Human database (downloaded 2021-01-14; 178486 sequences). Peptide precursor mass tolerance was set at 10 ppm and MS/MS tolerance at 0.6 Da. Search criteria included oxidation of methionine (+15.995 Da), acetylation of the protein N-terminus (+42.011 Da) and Methionine loss plus acetylation of the protein N-terminus (−89.03 Da) as variable modifications and carbamidomethylation of cysteine (+57.021 Da) and the addition of the TMT mass tag (+229.163 Da) to peptide N-termini and lysine as fixed modifications. Searches were performed with full tryptic digestion and a maximum of 2 missed cleavages were allowed. The reverse database search option was enabled and all data was filtered to satisfy false discovery rate (FDR) of 5%. Where proteins were identified and quantified by an identical group of peptides as the master protein of their protein group, these are designated ‘candidate master proteins’. Next, we used the annotation metrics for candidate master proteins retrieved from Uniprot to select the best annotated protein which was then designated as master protein. This enables us to infer biological trends more effectively in the dataset without any loss in the quality of identification or quantification.

For statistical analysis of differential protein abundance between conditions, standard t-test were used. Volcano plots were generated with VolcanoseR2 webapp (Goedhart & Luijsterburg, 2020) or GraphPad Prism 9 software (LaJolla, CA). Gene ontology analysis of the proteins identified was performed using Metascape webapp (Zhou et al., 2019) and Cytoscape 3.9 software to represent pathway enrichment, DisGeneNET category enrichment, and protein-protein interaction (PPI) networks. Dot plot diagrams were generated using the platform prohits-viz.org (Knight et al., 2017) to better visualize the quantitative enrichment and statistical significance of each identified protein across all data sets.

### Peptide synthesis

The human DENND4C peptide (1189 – AKVVQREDVETGLDPLSL - 1206) and the L1204A mutant (1189 – AKVVQREDVETGLDPASL - 1206) were made by solid phase peptide synthesis in house, purified by reverse phase HPLC and purity assessed by mass spectrometry. VPS29 binding cyclic peptide RT-D1 (Ac-yIIDTPLGVFLSSLKRC-NH2) was synthesized and prepared as described previously (Chen et al., 2021).

### Recombinant protein expression and purification

All the bacteria constructs were expressed in BL21 (DE3) cells using the autoinduction method. In brief, the culture containing complex medium was grown for 4 hours at 37°C then overnight at 20°C before harvesting for purification. To obtain the Retromer trimer, GST-tagged Vps29 and Vps35 co-expressed cell pellet were mixed with His-Vps26A and resuspended in lysis buffer containing 50 mM Tris-HCl pH 7.5, 200 mM NaCl, 2 mM 2-Mercaptoethanol, 50 μg/ml benzamidine and DNase I before passed through a Constant System TS-Series cell disruptor. Cell pellet containing GST-tagged VPS29 or GST-tagged hTBC1D13 was resuspended and lysed in the same procedures as Retromer. In all cases, the soluble fractions clarified by centrifugation was loaded onto the pre-equilibrated glutathione sepharose (GE healthcare) for initial purification. For Retromer, an additional Talon® resin purification was carried out to obtain the correct stoichiometry. GST-tag removal was performed co-column overnight by adding thrombin for Human Retromer and PreScission protease for TBC1D13. GST-tag removed fraction was further purified by size-exclusion chromatography (SEC) using a Superdex 200 (16/600) column (GE Healthcare) equilibrated with a buffer containing 50 mM Tris-HCl pH 7.5, 200 mM NaCl, 2 mM β-ME.

### Isothermal titration calorimetry (ITC)

The binding affinities between Retromer, DENND4C and TBC1D13 were determined using a Microcal PEAQ-ITC (Malvern) at 25°C. For Retromer and DENND4C, 1 mM of native or L1204A mutant DENND4C_1189-1206_ peptide was titrated into 14 µM Retromer. The effect of cyclic peptide was performed by pre-mixing 140 µM RT-D1 into 14 µM Retromer before titrating with 1 mM of native DENND4C_1189-1206_ peptide. For the binding experiment between Retromer and TBC1D13, 530 µM of TBC1D13 was titrated into 14 µM Retromer. The effect of the cyclic peptide was carried using the same concentration of RT-D1 as described above. The ITC running procedures consisted of an initial 0.4 μl (not used in data processing) followed by 12 serial injections of 3.22 μl each with 180 sec intervals. The resulting titration data were integrated with Malvern software package by fitting and normalized data to a single-site binding model, yielding the thermodynamic parameters for all binding experiments including dissociation constant (Kd), enthalpy (ΔH), Gibbs free energy (ΔG) and -TΔS. The stoichiometry (N) was refined initially, and if the value was close to 1, then it was set to exactly 1.0 for calculation. Data presented are the mean of triplicate titrations for each experiment.

### Crystallization and structure determination

Crystal of VPS29 – DENND4C_1189-1206_ peptide complex was grown by hanging drop vapor diffusion method at 20°C in the condition containing 0.1 M Tris-HCl pH 8.5, 0.2 M MgCl_2_, 30% PEG4000. Best quality crystal was obtained by adding 3 molar excesses of DENND4C_1189-1206_ peptide into the purified mVps29 to a final concentration of 12 mg/ml with a protein-reservoir drop ratio of 2:3. Crystals grew after 10 days were cryoprotected in reservoir solution with additional 10% glycerol. X-ray diffraction data were carried out at the Australian Synchrotron MX2 beamline at 100 K. Data collected from the Australian Synchrotron were indexed and integrated by AutoXDS and scaled using Aimless (Evans and Murshudov, 2013). Phase was solved by molecular replacement using Phaser (McCoy et al., 2007) with native mVps29 structure as the initial mode template. After refinement, the electron density corresponds to DENND4C was clearly visible. The refinement was performed using Phenix (Adams et al., 2010) with inspection of resulting model in Coot guided by F_o_ – F_c_ difference map. Molprobity (Chen et al., 2010) was used to evaluate the quality geometry of the refined and molecular figures were generated using PyMOL. Data collection and refinement statistics are summarized in Table 3. The VPS29 – DENND4C_1189-1206_ structure has been deposited at PDB with identification code 8VOD.

**Table 3.**
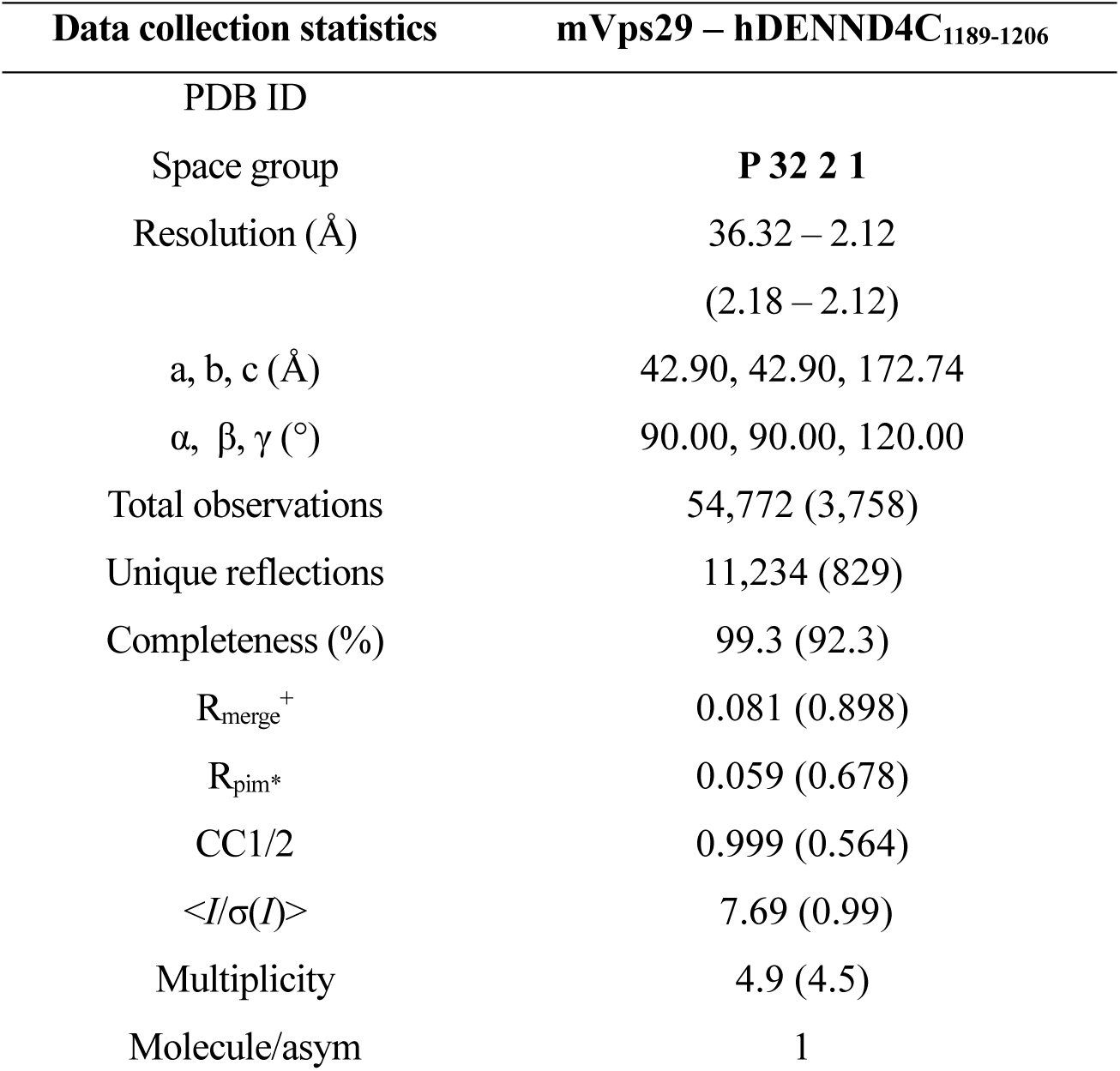
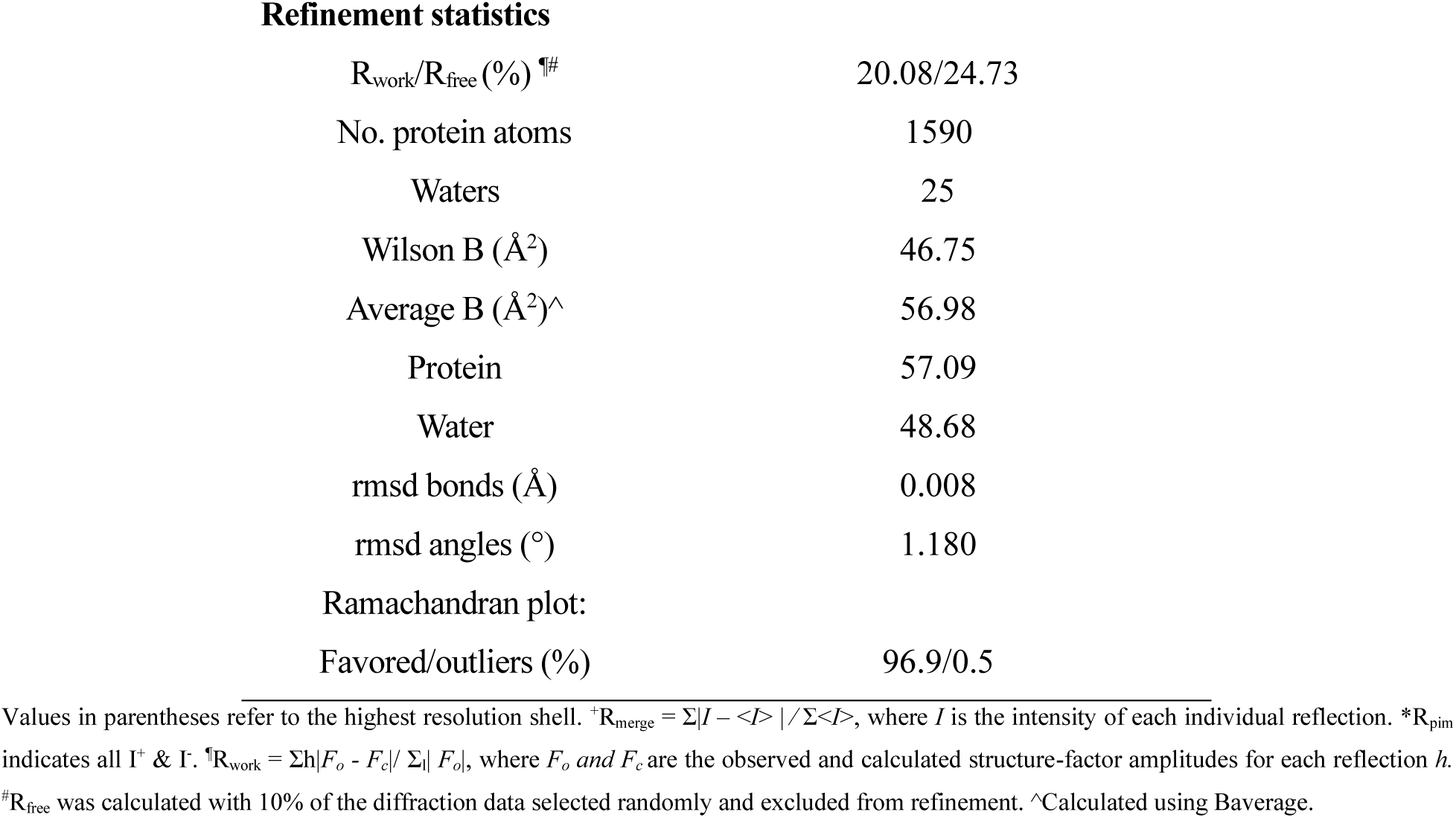
Summary of crystallographic structure determination statistics.

| Data collection statistics | mVps29 – hDENND4C<sub>1189-1206</sub> |
| --- | --- |
| PDB ID |  |
| Space group | **P 32 2 1** |
| Resolution ($\text{\AA}$) | 36.32 – 2.12<br>(2.18 – 2.12) |
| a, b, c ($\text{\AA}$) | 42.90, 42.90, 172.74 |
| $\alpha$, $\beta$, $\gamma$ ($^\circ$) | 90.00, 90.00, 120.00 |
| Total observations | 54,772 (3,758) |
| Unique reflections | 11,234 (829) |
| Completeness (%) | 99.3 (92.3) |
| $R_{\text{merge}}^+$ | 0.081 (0.898) |
| $R_{\text{pim}}^*$ | 0.059 (0.678) |
| CC1/2 | 0.999 (0.564) |
| $\langle I/\sigma(I) \rangle$ | 7.69 (0.99) |
| Multiplicity | 4.9 (4.5) |
| Molecule/asym | 1 |
| Refinement statistics |  |
| $R_{\text{work}}/R_{\text{free}} (\%)$ <sup>¶</sup> | 20.08/24.73 |
| No. protein atoms | 1590 |
| Waters | 25 |
| Wilson B ($\text{\AA}^2$) | 46.75 |
| Average B ($\text{\AA}^2$)<sup>^</sup> | 56.98 |
| Protein | 57.09 |
| Water | 48.68 |
| rmsd bonds ($\text{\AA}$) | 0.008 |
| rmsd angles ($^\circ$) | 1.180 |
| Ramachandran plot: |  |
| Favored/outliers (%) | 96.9/0.5 |

Values in parentheses refer to the highest resolution shell. <sup>\*</sup> $R_{\text{merge}} = \sum |I - \langle I \rangle| / \sum \langle I \rangle$ , where $I$ is the intensity of each individual reflection. <sup>\*</sup> $R_{\text{pim}}$ indicates all I<sup>+</sup> & I<sup>-</sup>. <sup>¶</sup> $R_{\text{work}} = \sum |F_o - F_c| / \sum |F_o|$ , where $F_o$ and $F_c$ are the observed and calculated structure-factor amplitudes for each reflection $h$ . <sup>#</sup> $R_{\text{free}}$ was calculated with 10% of the diffraction data selected randomly and excluded from refinement. <sup>^</sup>Calculated using Baverage.

### Protein structural prediction, modelling and visualisation

All protein models were generated using AlphaFold2 Multimer implemented in the Colabfold interface available on the Google Colab platform. Due to the protein length, the Alphafold3 was used to screen for potential interaction between full-length Retromer and TBCs/DENNDs. In the case of Alphafold2 modelling experiment, ColabFold was executed using default settings where multiple sequence alignments were generated with MMseqs2. For all final models displayed in this manuscript, structural relaxation of peptide geometry was performed with AMBER. For all modelling experiments, we assessed (i) the prediction confidence measures (pLDDT and interfacial iPTM scores), (ii) the plots of the predicted alignment errors (PAE) and (iii) backbone alignments of the final structures. The confidence level of Retromer with TBCs/ DENNDs were calculated using the sum of IPTM score generated by Alphafold3 (full-length Retromer) and Alphafold2 (sub-complex and individual subunits). For clarity, the confidence level score was differentiated into two groups depending on the Retromer subcomplex (Vps26A – Vps35 and Vps29 – Vps35). SPOC score was calculated using the web-based tool (https://predictomes.org/tools/spoc/) using default setting described previously. Similar, pDOCK score was obtained using previously established method. All structural images were made with Pymol (Schrodinger, USA; https://pymol.org/2/).

### Statistical analysis

Statistics from western blots and confocal microscopy from a minimum of 3 independent experimental repeats was generated and represented using GraphPad Prism 9 software (LaJolla, CA). Graphs were plotted representing the mean value ± the standard deviation (SD) for each experimental condition. n represents the number of independent experimental repeats. In all graphs, * = p < 0.05, ** = p < 0.01, *** = p < 0.001, **** = p < 0.0001.

## Supporting information

Excel Spreadsheet Proximity Proteins

## ACKNOWLEDGEMENTS

We thank the Wolfson Bioimaging Facility at the University of Bristol for their support. Work in the Cullen laboratory is supported by the Wellcome Trust (104568/Z/14/Z and 220260/Z/20/Z), the Medical Research Council (MR/L007363/1 and MR/P018807/1), the Lister Institute of Preventive Medicine, and the award of a Royal Society Noreen Murray Research Professorship to P.J.C. (RSRP/R1/211004). We acknowledge use of the University of Queensland Remote Operation Crystallization and X-ray (UQ ROCX) Facility and the assistance of K. A. Arachchige. X-ray data were collected on the MX2 beamline at the Australian Synchrotron. B.M.C. is supported by an Investigator Grant from the National Health and Medical Research Council (APP2016410). C.A.P was initially supported by Beca Fundación Ramón Areces Estudios Postdoctorales en el Extranjero.

## AUTHOR CONTRIBUTIONS

BioID and cell biology analysis: C.A-P. and A.J.E. Biophysics, structural analysis and AlphaFold2: K.E.C., Q.C. and M.L. Proteomics and Bioinformatic analysis: P.A.L. and K.J.H. Manuscript Writing - 1^st^ draft: C.A-P., K.E.C., B.M.C. and P.J.C; Final Version: all authors. Initial Concept: C.A-P. and P.J.C. Concept Development: all authors. Funding and Supervision: K.A.W., B.M.C. and P.J.C.

## CONFLICTS OF INTEREST

The authors declare that they have no conflict of interest.

---

**Supplementary Figure 1.**
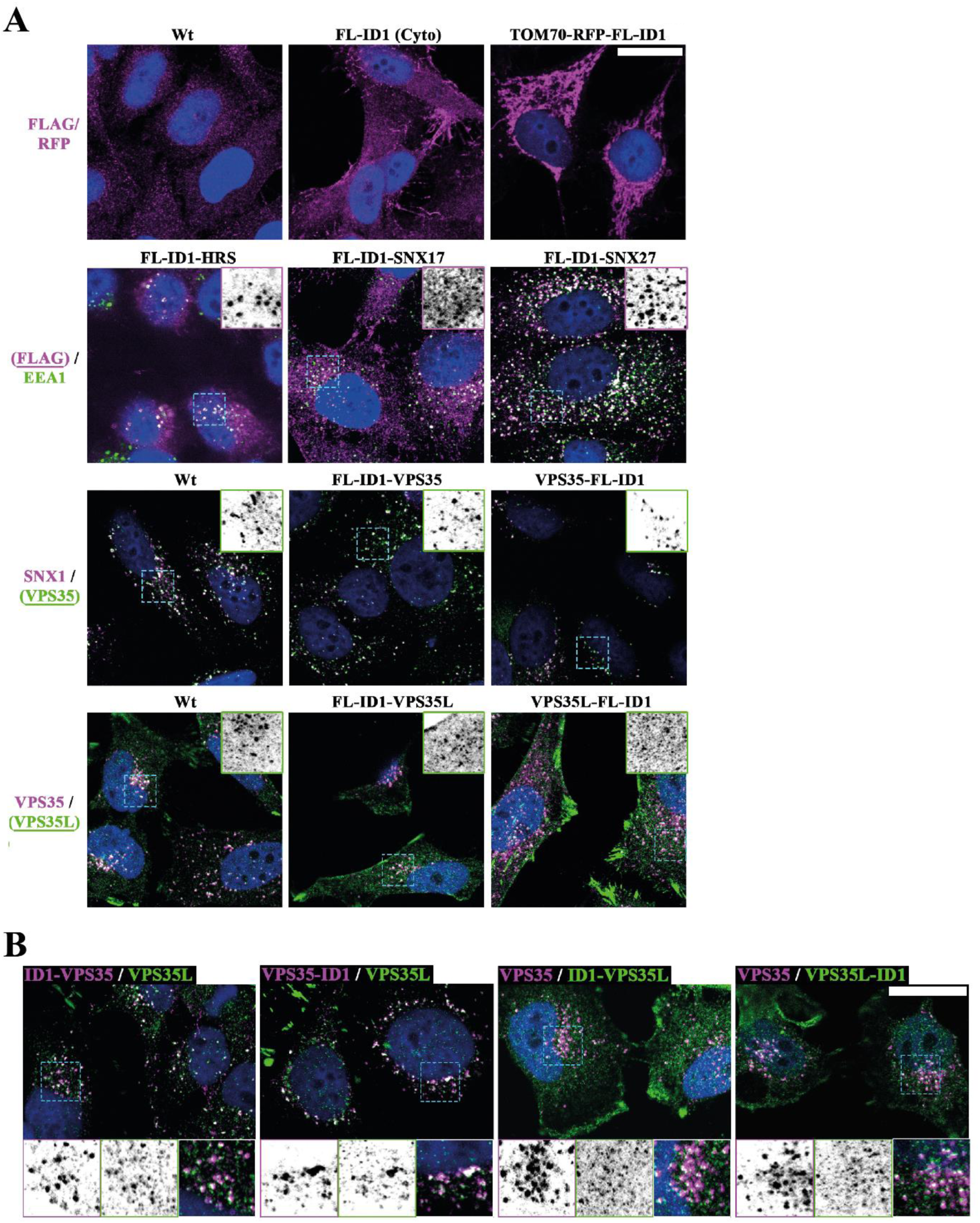
Validation of localization of BioID1-tagged proteins. **(A)** Confocal images showing validation of localization of BioID1-tagged proteins/locations using specific antibodies or anti-FLAG. The FLAG epitope is coded just before the BioID1 sequence and ‘FL-ID1’ is abbreviation for FLAG epitope followed by BioID1 biotin ligase. **(B)** Confocal images showing colocalization between Retromer (VPS35) and Retriever (VPS35L) complexes in cells expressing the different BioID1-tagged versions of VPS35 or VPS35L. Scale bar - 20 μm.

**Supplementary Figure 2.**
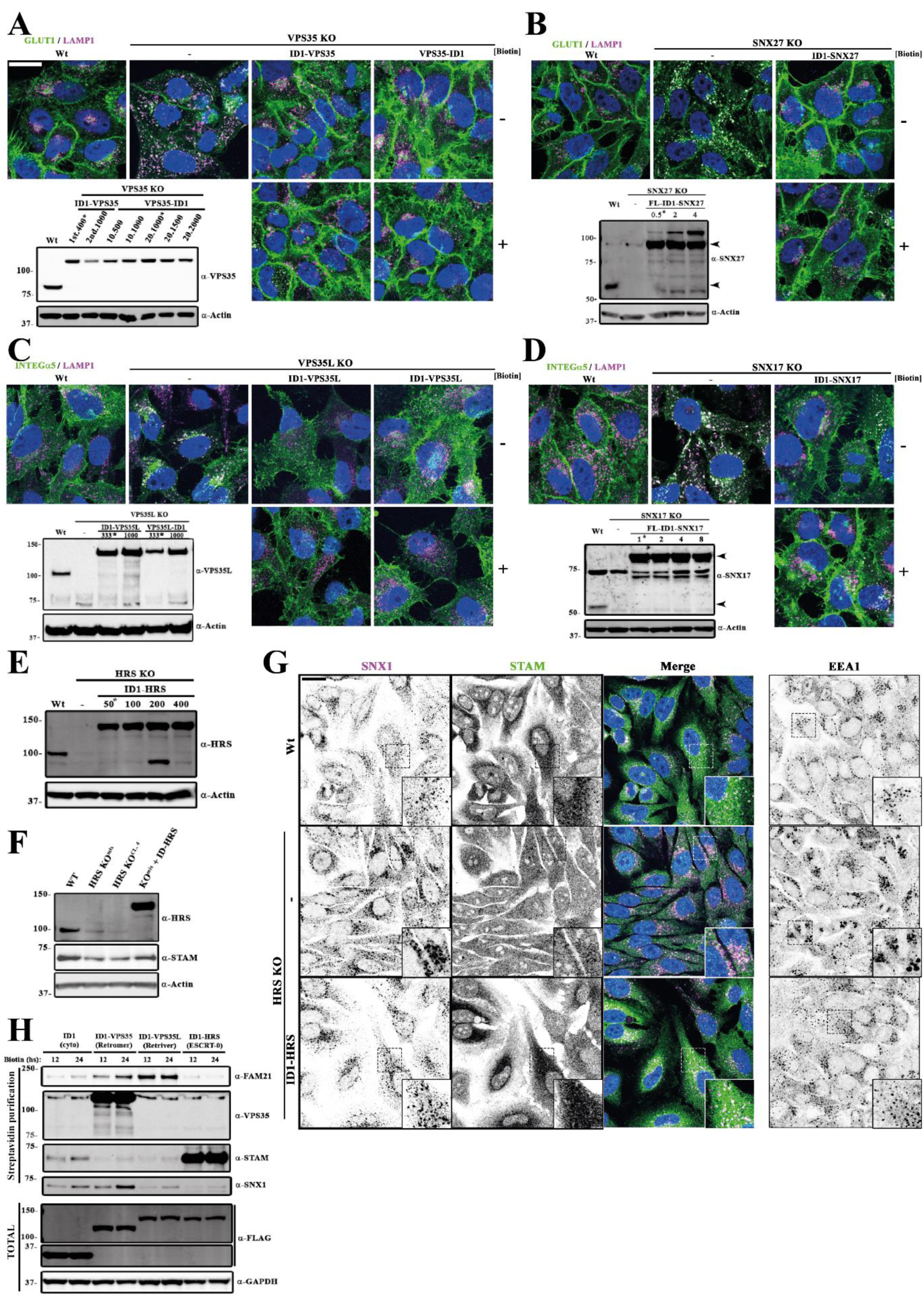
Validation of functionality of BioID1-tagged proteins. **(A)** Validation of BioID1 Retromer (VPS35) engineered cells lines; ID1-VPS35 and VPS35-ID1. Previously characterized HeLa VPS35 KO cell line was transduced with lentivirus to express ID1-VPS35 or VPS35-ID1. By means of viral titration and puromycin selection populations with VPS35 levels comparable to the endogenous were selected. In the left bottom western blot (WB), asterisk indicates cell populations used for later studies. The functionality of the chimeras ID1-VPS35 and VPS35-ID1 was confirmed by analyzing the rescue of GLUT1 lysosomal missorting observed in HeLa VPS35 KO cells. The confocal images show colocalization of GLUT1 with lysosomal marker LAMP1. GLUT1 recovers its plasma membrane localization when VPS35 BioID1 versions are expressed, at basal conditions (top row images) and after 24 hrs incubation with 50 mM biotin (bottom row images). **(B)** Validation of ID1-SNX27 engineered cell line. Previously characterized HeLa SNX27 KO cell line was transduced with lentivirus to express ID1-SNX27 and by means of viral titration and puromycin selection we isolated population with SNX2 7 levels comparable to the endogenous. In the left bottom WB, asterisk indicated cell population used for later studies. The functionality of the chimera ID1-SNX27 was confirmed by analyzing the rescue of GLUT1 lysosomal missorting observed in HeLa SNX27 KO cells, following the same procedure as in (A). **(C)** Validation of BioID1 Retriever (VPS35L) engineered cells lines; ID1-VPS35L and VPS35L-ID1. Previously characterized HeLa VPS35L KO cell line was transduced with lentivirus to express ID1-VPS35L or VPS35L-ID1 and by means of viral titration and puromycin selection we isolated populations with levels comparable to the endogenous VPS35L (left bottom WB, asterisk indicated cell populations used for later studies). The functionality of the chimeras ID1-VPS35L and VPS35L-ID1 was confirmed by analyzing the rescue of α5-integrin lysosomal missorting observed in HeLa VPS35L KO cells. The confocal images show colocalization of α5-integrin with lysosomal marker LAMP1. α5-integrin recovers its plasma membrane localization when VPS35L BioID1 versions are expressed, at basal conditions (top row images) and after 24 hrs incubation with 50 mM biotin (bottom row images). **(D)** Validation of ID1-SNX17 engineered cell line. Previously characterized HeLa SNX17 KO cell line was transduced with lentivirus to express ID1-SNX17 and by means of viral titration and puromycin selection we isolated populations with the closest possible levels compared to endogenous SNX17 (left bottom WB, asterisk indicated cell population used for later studies). The functionality of the chimera ID1-SNX17 was confirmed by analyzing the rescue of α5-integrin lysosomal missorting observed in HeLa SNX17 KO cells, following the same procedure as in (C). **(E-G)** Validation of ID1-HRS engineered cell line. For the generation of HeLa CRISPR KO for HRS/HGS, we use a guide against HRS/HGS exon 3 (Table S1). **(E)** Single clonal HRS KO cell line was transduced with lentivirus to express ID1-HRS and by means of viral titration and puromycin selection we isolated populations with the closest possible levels compared to endogenous HRS/HGS; asterisk indicated cell population used for later studies. **(F)** Western blot showing the rescue of total levels of STAM1/2 protein when ID1-HRS version is expressed. HRS/HGS and STAM1/2 form the ESCRT-0 complex. **(G)** Confocal images showing the rescue of SNX1 endosome morphology, STAM1/2 localization (left panels) and morphology of early endosomal marker EEA1 (right panels) when ID1-HRS version is expressed. **(H)** Western blot showing comparison of the biotinylated proteins after 12- or 24-hours incubation with 50 mM biotin, lysis and streptavidin purification among different cell lines. Top part shows biotinylated proteins, and bottom part shows total protein levels. Scale bar - 20 μm.

**Supplementary Figure 3.**
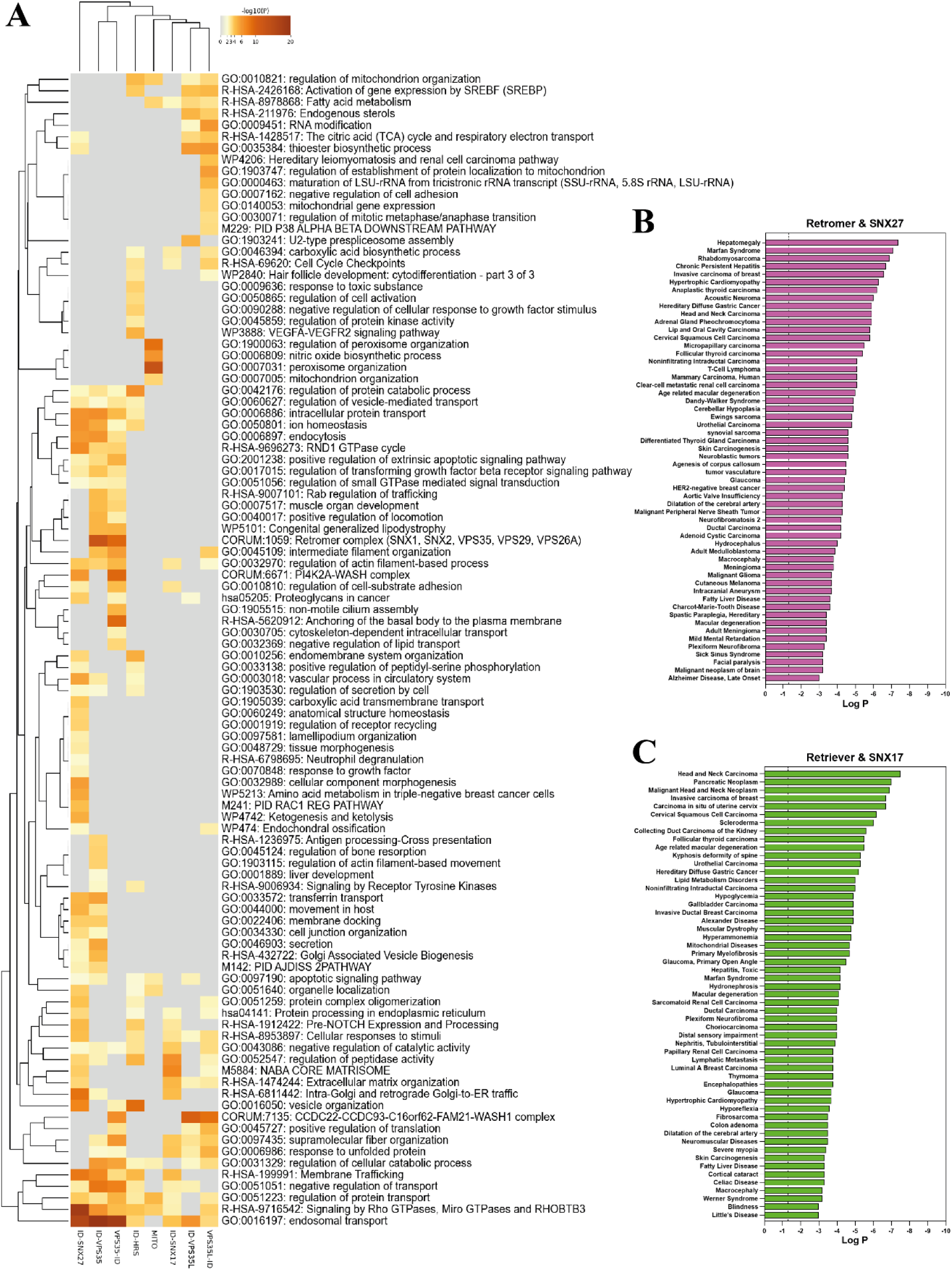
Gene ontology (GO) comparison among all proximity proteomes; extended versions. **(A)** Heat map representing the comparison of the main GO terms associated to each proximity proteome. The colour gradient represents the enrichment significance for each GO term (-Log_10_); for reference 1.3 −Log_10_ is equal to 0.05 p-value. **(B-C)** Disease associated GO terms and its significance (-Log_10_) for Retromer and SNX27 proximity proteomes combined (B), and for the combined Retriever and SNX17 proximity proteomes (C).

**Supplementary Figure 4.**
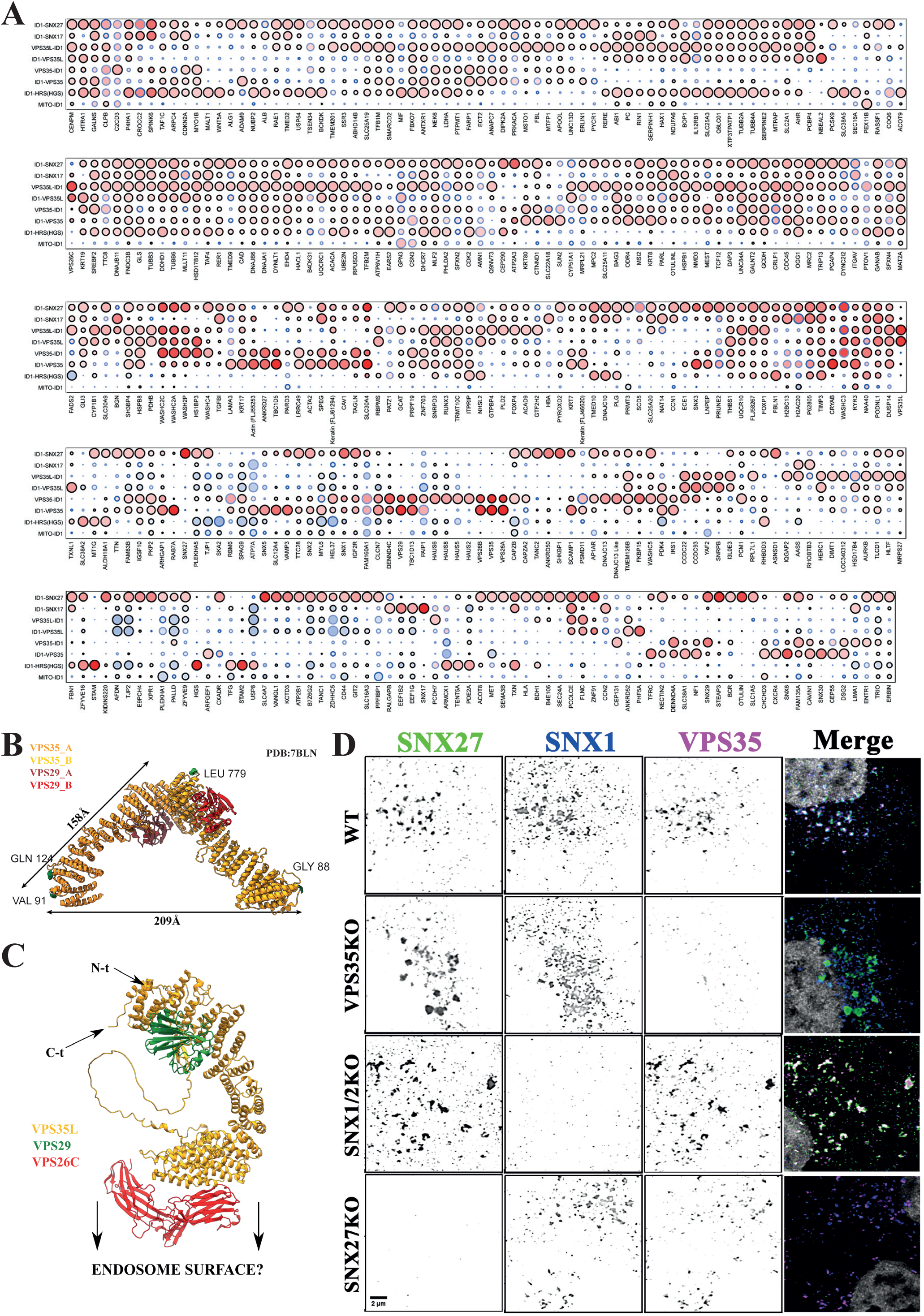
Proteomic quantitative comparison among endosomal sub-domains; extended version. **(A)** Dot plot showing the comparison of FC (Log_2_) and significance of detected proteins, along the different endosomal sub-domains as well Mito-BioID1. Only proteins with FC ≥ 0.8 and significant (p-value ≤ 0.05) in at least one proximity proteome were selected for the comparison. Filling colours represent the FC value, edge colours the p-value and the size symbolize the relative FC along the different sub-domains for that specific protein (see dot-plot legend in Figure 2E). **(B)** Representation of main length distances over the published structure of VPS35/VPS29 arch from the metazoan membrane-assembled Retromer:SNX3 complex modelled with human proteins (PDB:7BLN). **(C)** AlphaFold2 predictive model of Retriever complex. As Retriever complex binds the cargo adaptor SNX17 through its VPS26C subunit in the vicinity of the endosomal surface, the N and C termini of VPS35L might points far from the endosomal surface. This could explain why both Retriever BioID1 versions, ID1-VPS35L and VPS35L-ID1, strongly biotinylate WASH subunits and CCDC22/93, but not SNX17 cargo adaptor. **(D)** Images showing colocalization of Retromer (VPS35), SNX27 and SNX1 in HeLa parental, VPS35KO, SNX1/SNX2 double KO, and SNX27KO using endogenous antibodies and confocal imaging. Images were acquired on a Leica SP8 multi-laser point scanning confocal microscope with a 63x NA1.4 UV oil-immersion lens and using the Leica ‘Lightning’ mode for adaptive deconvolution to improve lateral resolution. Scale bar represents 2 μm. Images from top wild-type panel of this figure are shown in Figure 2I.

**Supplementary Figure 5.**
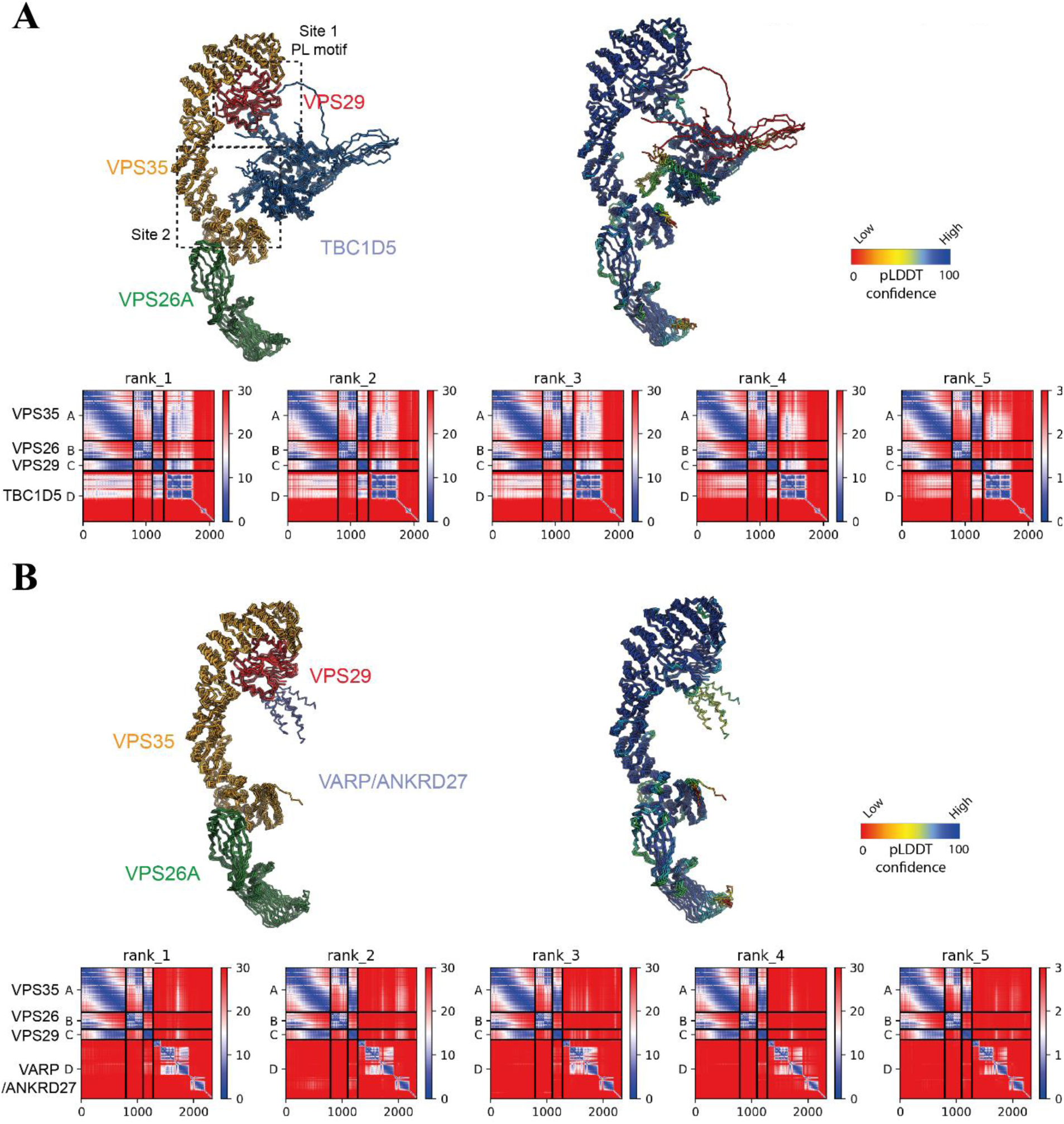
Retromer interactions with TBC1D5 and VARP/ANKRD27. **(A)** AlphaFold2-predicted complex between human Retromer and TBC1D5 (Q92609). The top five ranked structures are aligned and shown in ribbon diagram. For clarity, the extended C-terminal disordered sequences of TBC1D5 are omitted from the images. The Left panel shows each chain coloured with VPS35 (orange), VPS26A (green), VPS29 (red) and TBC1D5 (blue). The right panel shows the structures coloured according to the pLDDT confidence score. Bottom panels show the predicted alignment error (PAE) plots for all five predictions of each complex. **(B)** AlphaFold2-predicted complex between human Retromer and VARP/ANKRD27 (Q96NW4). The top five ranked structures are aligned and shown in ribbon diagram. For clarity, only the primary binding regions of VARP are shown in the images. The Left panel shows each chain coloured with VPS35 (orange), VPS26A (green), VPS29 (red) and VARP (blue). The right panels show the structures coloured according to the pLDDT confidence score. Bottom panels show the PAE plots for all five predictions of each complex.

**Supplementary Figure 6.**
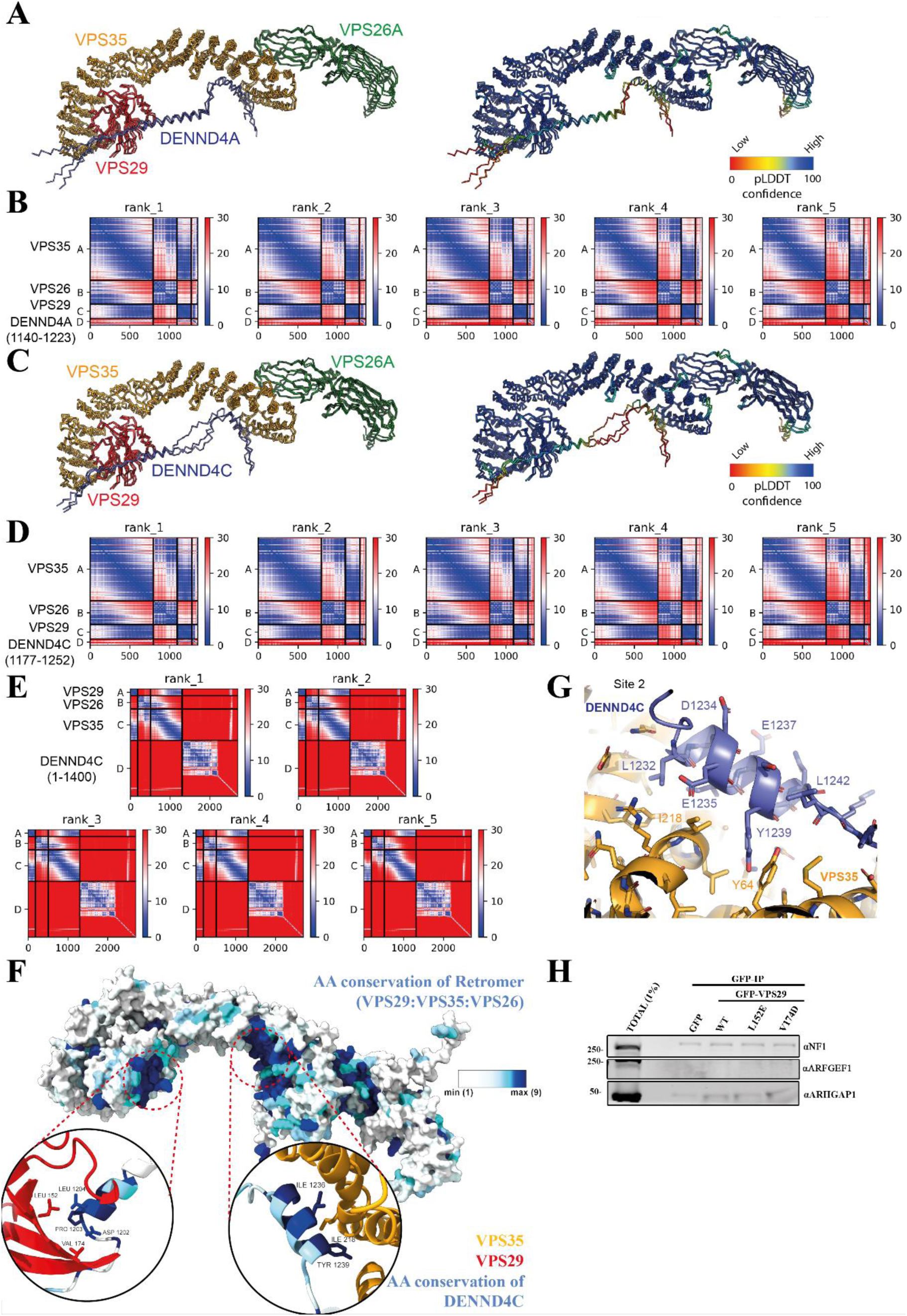
Retromer interactions with DENND4A and DENND4C. **(A-D)** AlphaFold2-predicted complex between human Retromer and the binding regions of DENND4A residues 1140-1223 (Q7Z401) and DENND4C residues 1177-1252 (Q5VZ89). The top three ranked structures are aligned and shown in ribbon diagram. (A and C) The Left panel shows each chain coloured with VPS35 (orange), VPS26A (green), VPS29 (red) and DENND4A/C (blue). The right panels show the structures coloured according to the pLDDT confidence score. (B and D) Graphics show the predicted alignment error (PAE) plots for all five predictions of each complex. **(E)** Graphics show the PAE plots for all five predictions of each complex between Retromer and first 1400 amino acids from DENND4C (Q5VZ89) predicted by AlphaFold2. **(F)** Amino acid sequence conservation depicted over AlphaFold2 models of Retromer complex and DENND4C interacting domains calculated with ConSurf server applying default parameters. Main structure corresponds to Retromer trimer (VPS26A:VPS29:VPS35) with the surface representation. Circular insets show magnifications for the 2 predicted binding sites between DENND4C and Retromer. Left inset shows the primary binding site consisting of a conserved PL motif (DENND4C ^1203^PL^1204^) interacting with the Leu152 containing hydrophobic cavity of VPS29. Right inset shows a second binding site predicted between a short DENND4C α-helical stretch (residues 1232-1243) with α-helices towards the amino-termini of VPS35. For clarity, ribbon representation was selected and only DENND4C sequence conservation at binding sites was shown. **(G)** Detailed view of Alphafold2 model of the secondary DENND4C binding site on the VPS35 subunit of Retromer. **(H)** GFP based co-immunoprecipitation (co-IP) of GFP-VPS29 wild-type (WT) and hydrophobic pocket mutants after transient transfection in HEK293T cells. Additional GTPase regulators found in the Retromer proximity proteomes (Figure 3C); neurofibromin (NF1), ARHGAP1 and ARFGEF1 show no interaction under GFP-VPS29 co-IP. Blots are from the same immunoprecipitation experiment as shown in Figure 6C

**Supplementary Figure 7.**
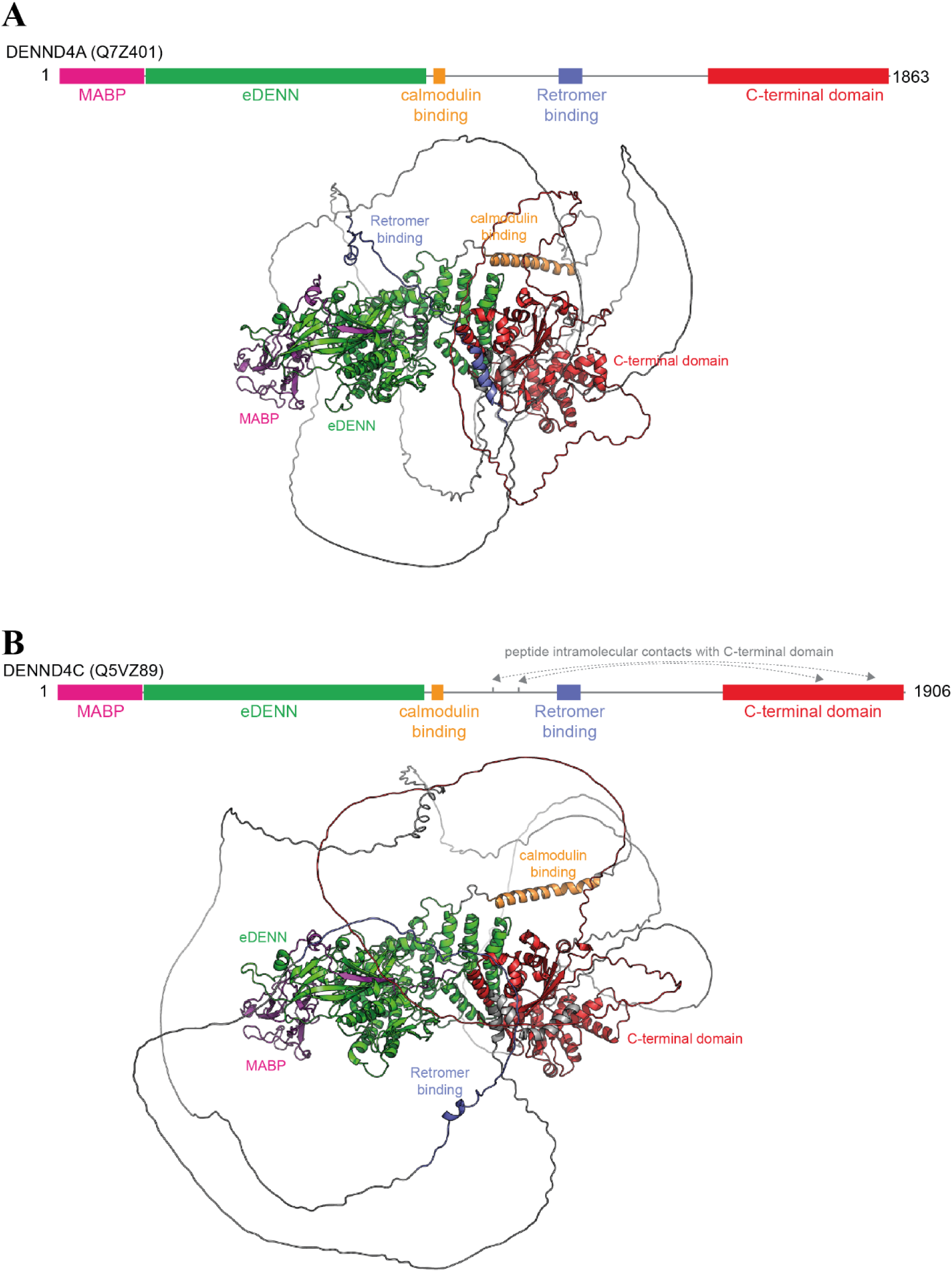
AlphaFold2 model of full length human DENND4A and DENND4C. **(A-B)** Top diagrams show domain architecture of human DENND4A (Q7Z401) (A) or DENND4C (Q5VZ89) (B) based on AlphaFold2 predictions of tertiary structures. Bottom diagrams show tertiary structure of DENND4A/ and DENND4C derived from AlphaFold2 prediction. Specific domains and binding sequences are coloured as indicated. MABP domain (MVB12-associated β-prism domain); eDENN domain (extended DENN domain); C-terminal domain (a globular domain that is structurally unique to DENND4 homologues).

**Supplementary Figure 8.**
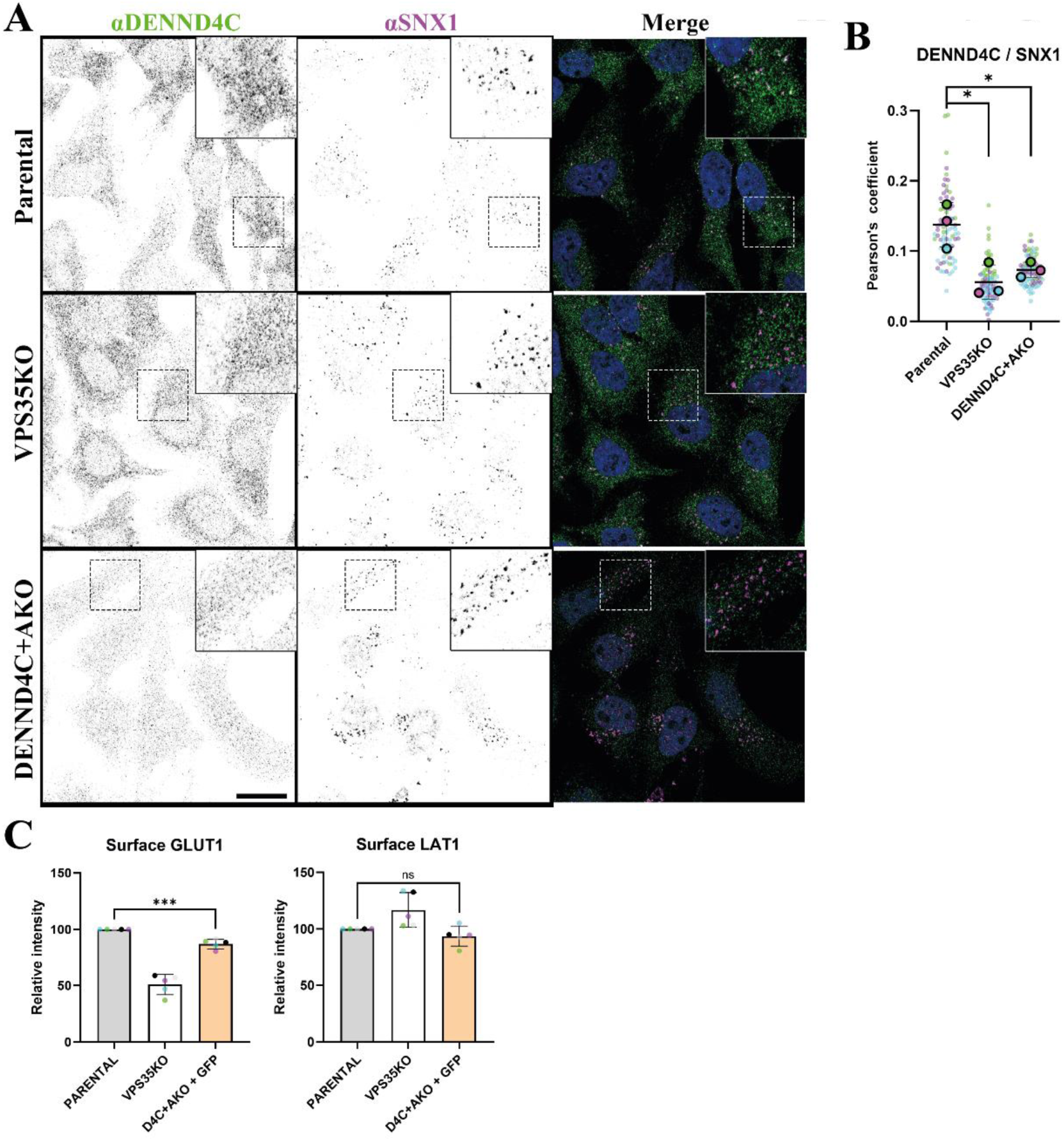
Extended figure on role of DENND4A/C in Retromer’s cell biology. **(A)** Confocal images showing colocalization of endogenous DENND4C and SNX1 in HeLa parental, VPS35KO and DENND4C and DENND4A double KO (DENND4C+A KO). Scale bar - 20 μm. **(B)** Quantitation and statistical analysis of colocalization between DENND4C and SNX1 using Pearson’s coefficient, showing a significant reduction in both KO lines. **(C)** Quantitation and statistical analysis of GLUT1 and LAT1 surface levels in Hela parental versus DENND4C+A KO from Figure 5C. n = 5 independent experiments. T-test analysis, data presented as mean values relative to WT and error bars represent SD. Data shows a moderate but significant reduction in GLUT1 surface levels.

**Supplementary Figure 9.**
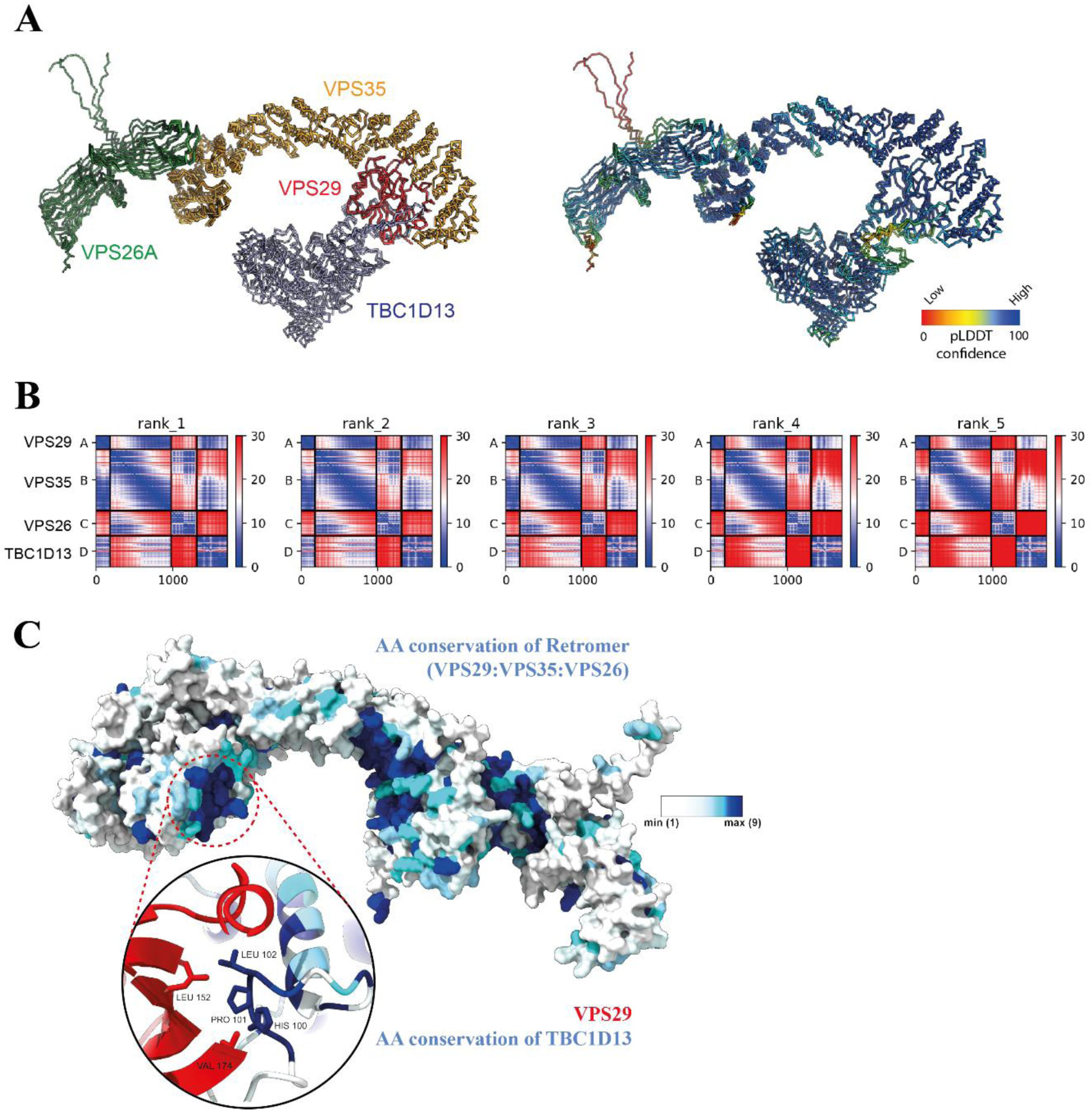
Retromer interactions with TBC1D13. **(A)** AlphaFold2-predicted complex between human Retromer and TBC1D13 (Q9NVG8). The top three ranked structures are aligned and shown in ribbon diagram. The Left panel shows each chain coloured with VPS35 (orange), VPS26A (green), VPS29 (red) and TBC1D13 (blue). The right panel show the structures coloured according to the pLDDT confidence score. **(B)** Predicted alignment error (PAE) plots for all five predictions from (A). **(C)** Amino acid sequence conservation depicted over AlphaFold2 model of Retromer complex and TBC1D13 interacting domains calculated with ConSurf server applying default parameters. Main structure corresponds to Retromer trimer with the surface representation. Circular inset shows magnification for the main binding site consisting of a conserved PL motif (TBC1D13 ^101^PL^102^) interacting with the Leu152 containing hydrophobic cavity of VPS29. For clarity, ribbon representation was selected and only TBC1D13 sequence conservation at binding site was shown.

**Supplementary Figure 10.**
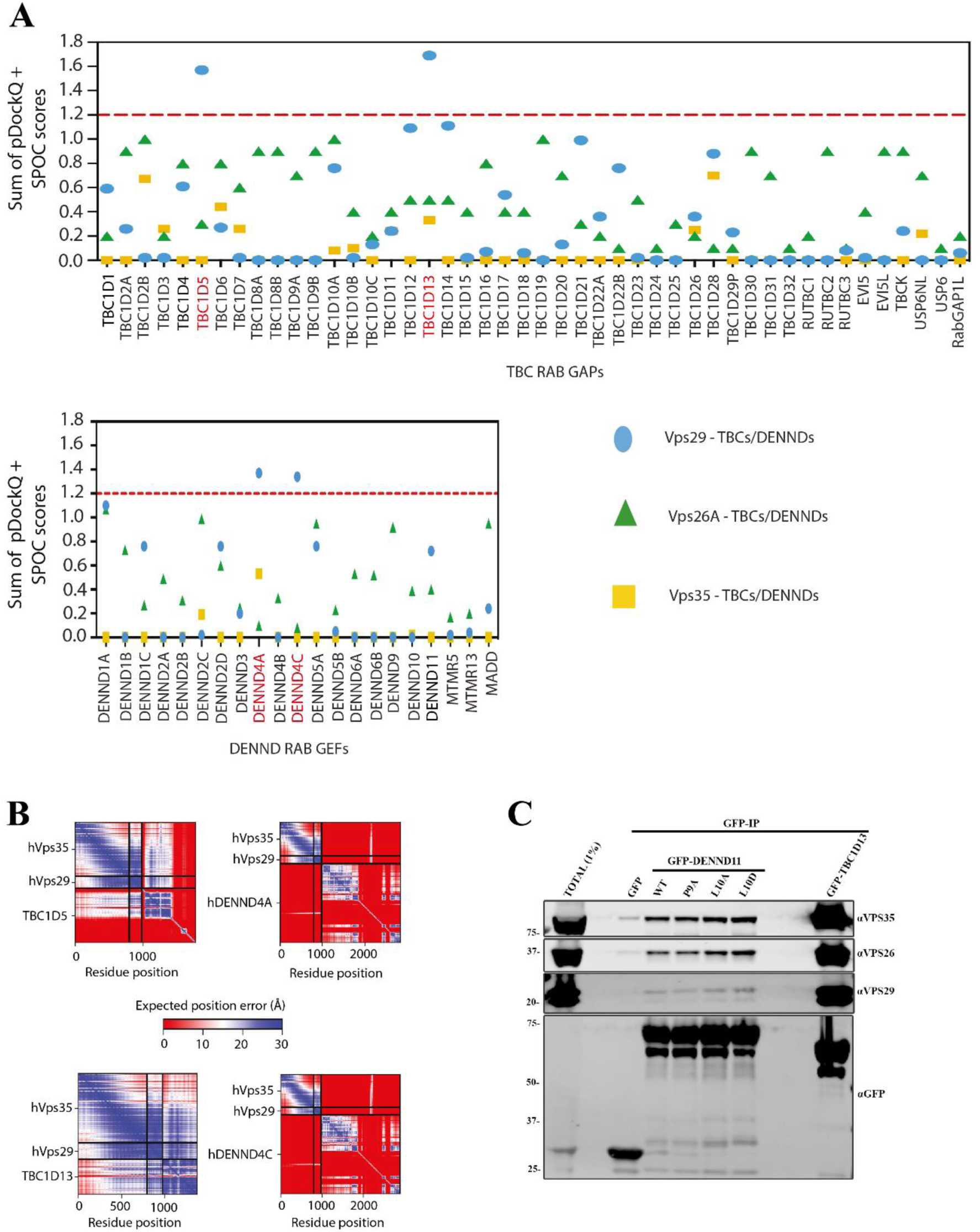
Retromer as a hub for RAB GTPase switch regulation: Extended information on VPS29 interaction screen. (**A**) Sum of pDockQ plus SPOC scores for all TBC and DENND domain containing RAB GAPs and GEFs respectively against VPS29, VPS26A and VPS35 subunits of human Retromer. **(B)** Top ranked PAE plots for VPS35:VPS29 dimer for their AlphaFold2 predicted association with TBC1D5, DENND4A, TBC1D13 and DENND4C. **(C)** GFP based co-immunoprecipitation (co-IP) of GFP-DENND11 wild-type (WT) and PL motive mutants compared to GFP-TBC1D13 after transient transfection in HEK293T cells.

**Supplementary Figure 11.**
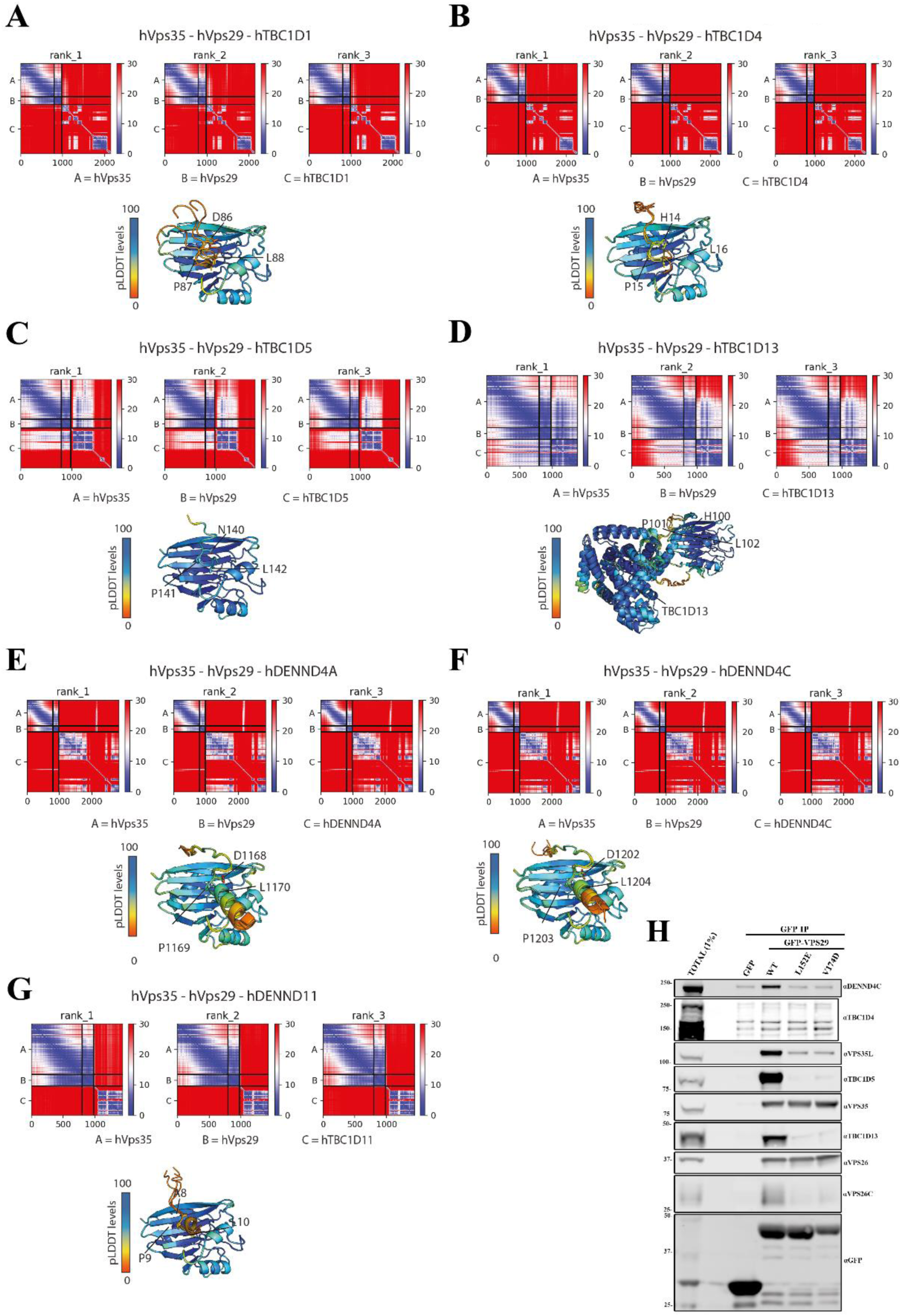
TBC1D1/4 interaction with Retromer. **(A-G)** Predicted alignment error (PAE) plots and pLDDT structural representation for all seven AlphaFold2 predictions of VPS35:VPS29 association with (A) TBC1D1, (B) TBC1D4, (C) TBC1D5, (D) TBC1D13, (E) DENND4A, (F) DENND4C, and (G) DENND11. **(H)** GFP based co-immunoprecipitation (co-IP) of GFP-VPS29 wild-type (WT) and hydrophobic pocket mutants after transient transfection in HEK293T cells. Here we blotted for TBC1D4, which contrary to DENND4A/C or TBC1D13 show weak interaction under GFP-VPS29 co-IP.

**Supplementary Figure 12.**
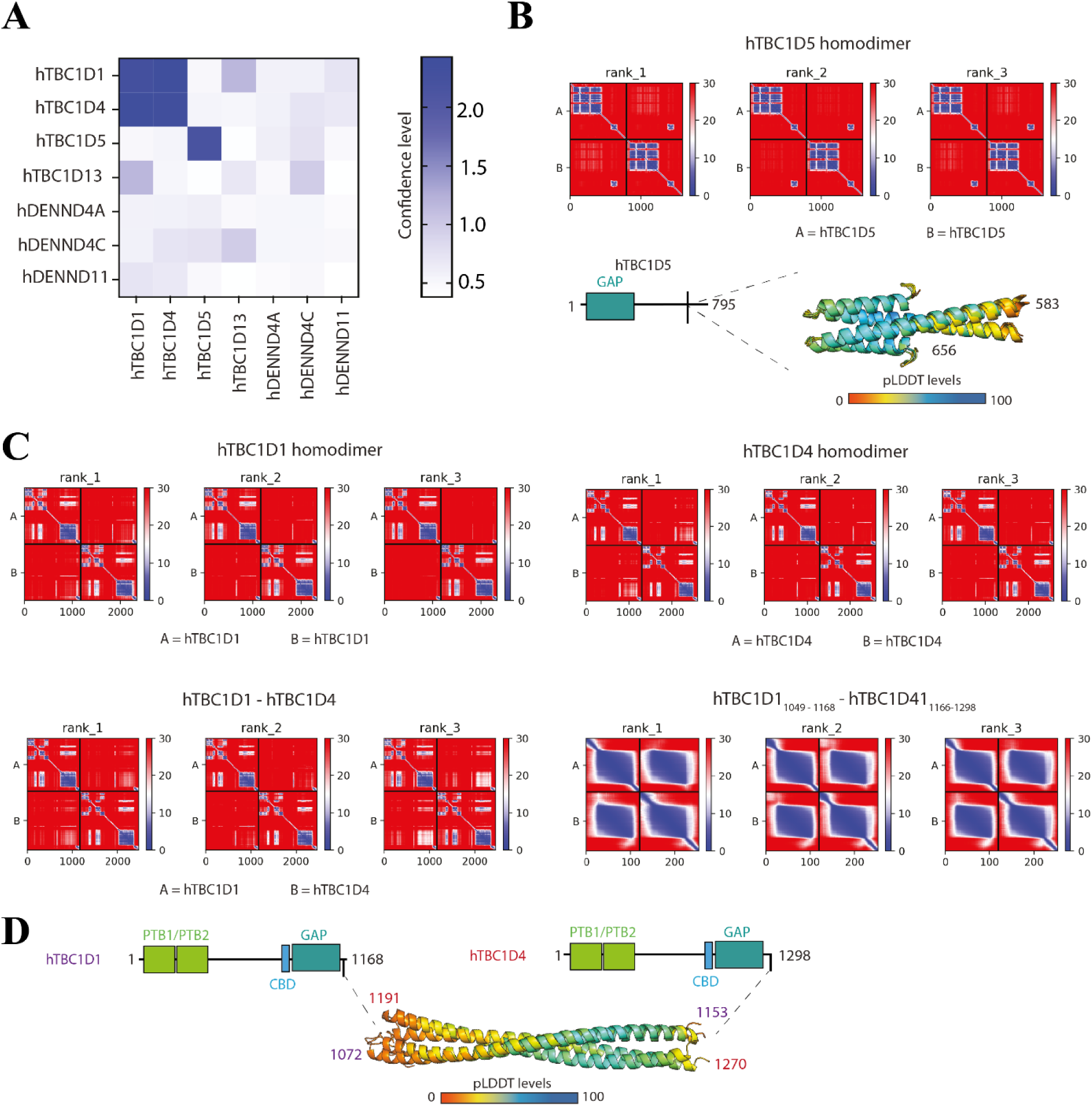
Predicted mechanism for TBC1D1-TBC1D4 heterodimer and TBC1D5 homodimer formation. **(A)** Confidence level predictor of homo- and heterodimer formation with a network of Retromer binding TBC and DENND proteins. **(B)** First three ranked PAE plots of the predicted formation of a TBC1D5 homodimer through carboxy-terminal coiled-coil interactions. **(C)** First three ranked PAE plots of the predicted formation of TBC1D1 homodimer, TBC1D4 homodimer, full length TBC1D1 and TBC1D4 heterodimer, and heterodimer of the carboxy-terminal regions of TBC1D1 and TBC1D4. **(D)** pLDDT levels depicted on the predicted AlphaFold model of the coiled-coil carboxy-terminal TBC1D1 and TBC1D4 heterodimer.

**Table S1:**
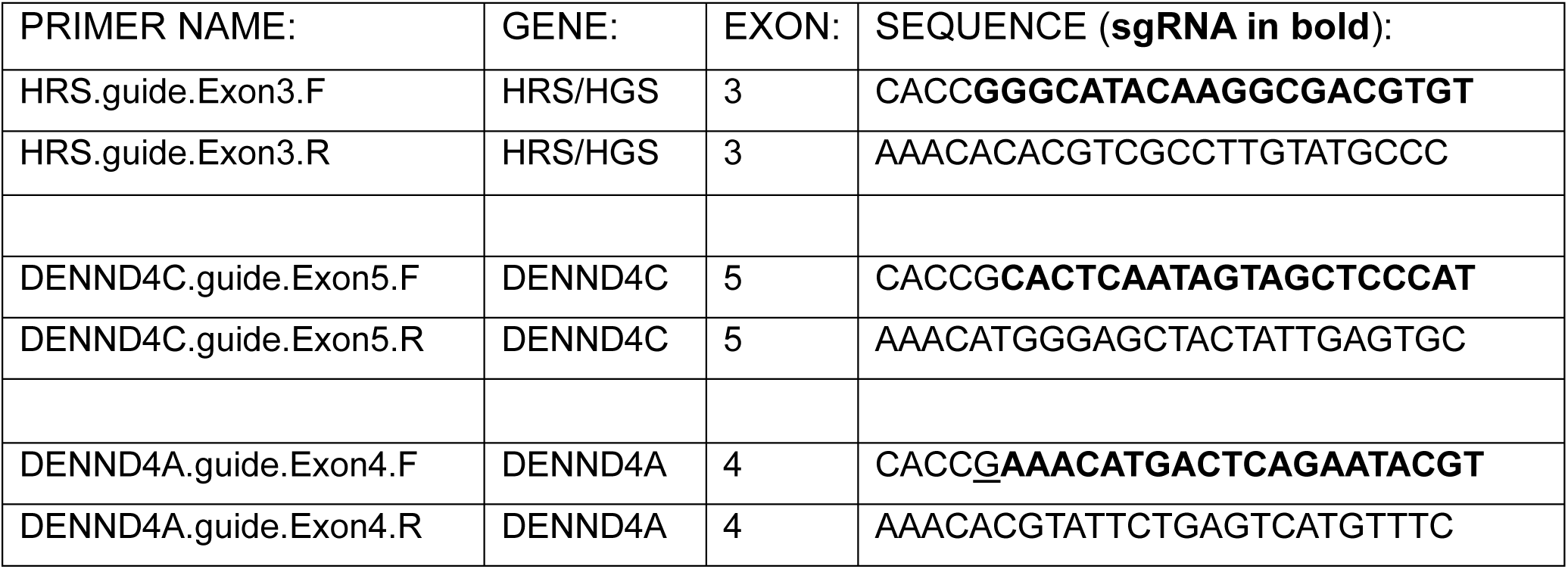
gRNAs.

**Table S2:**
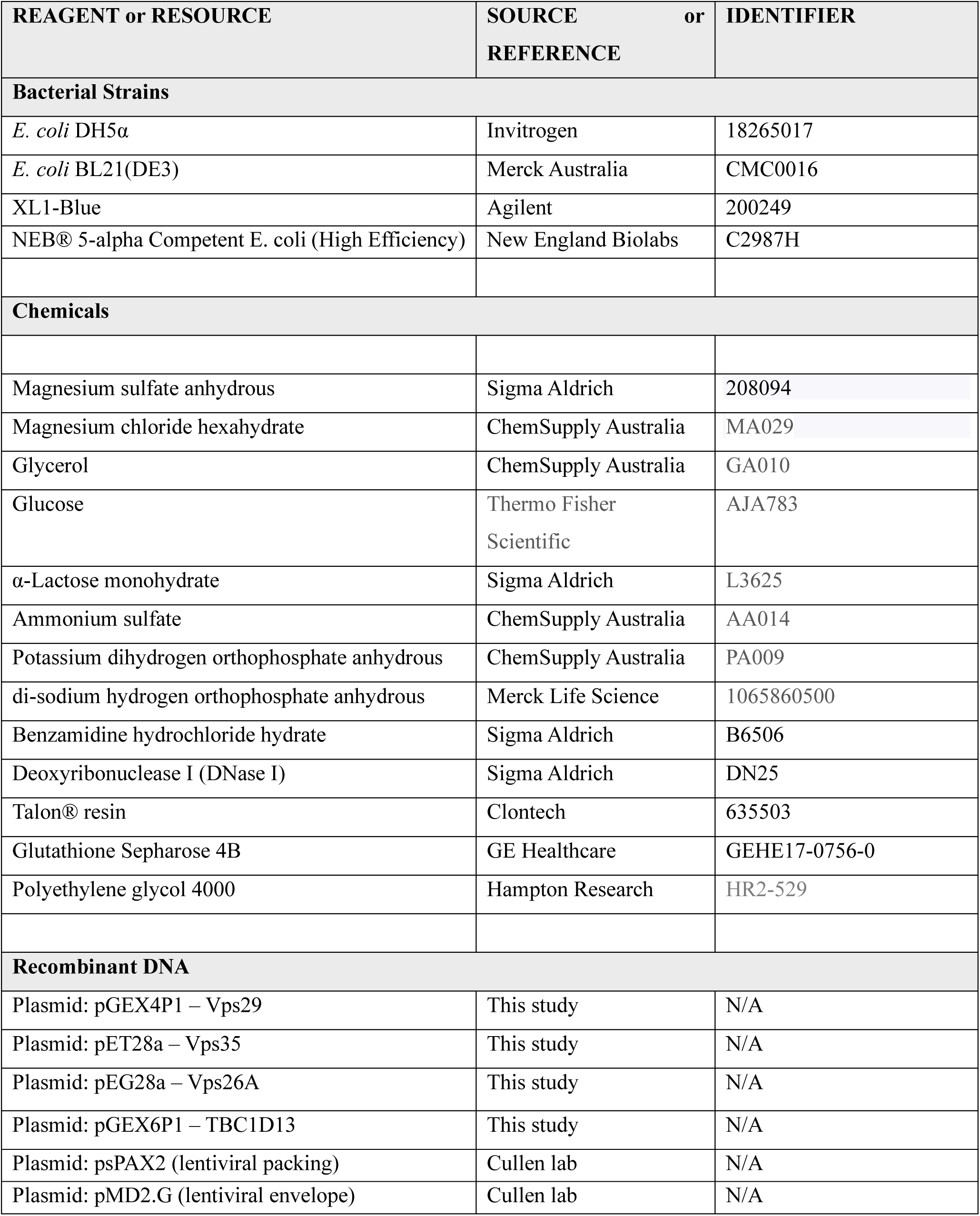
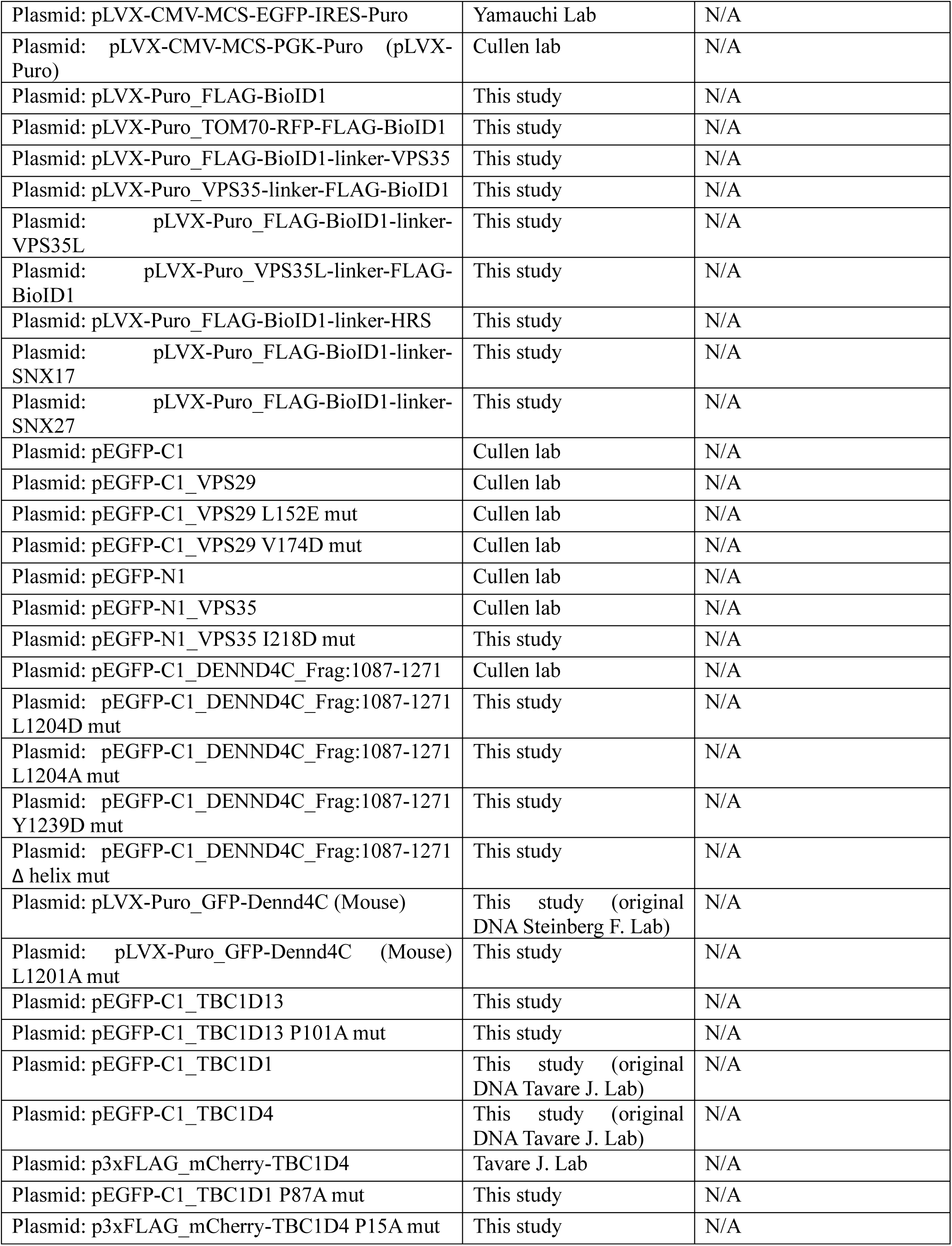
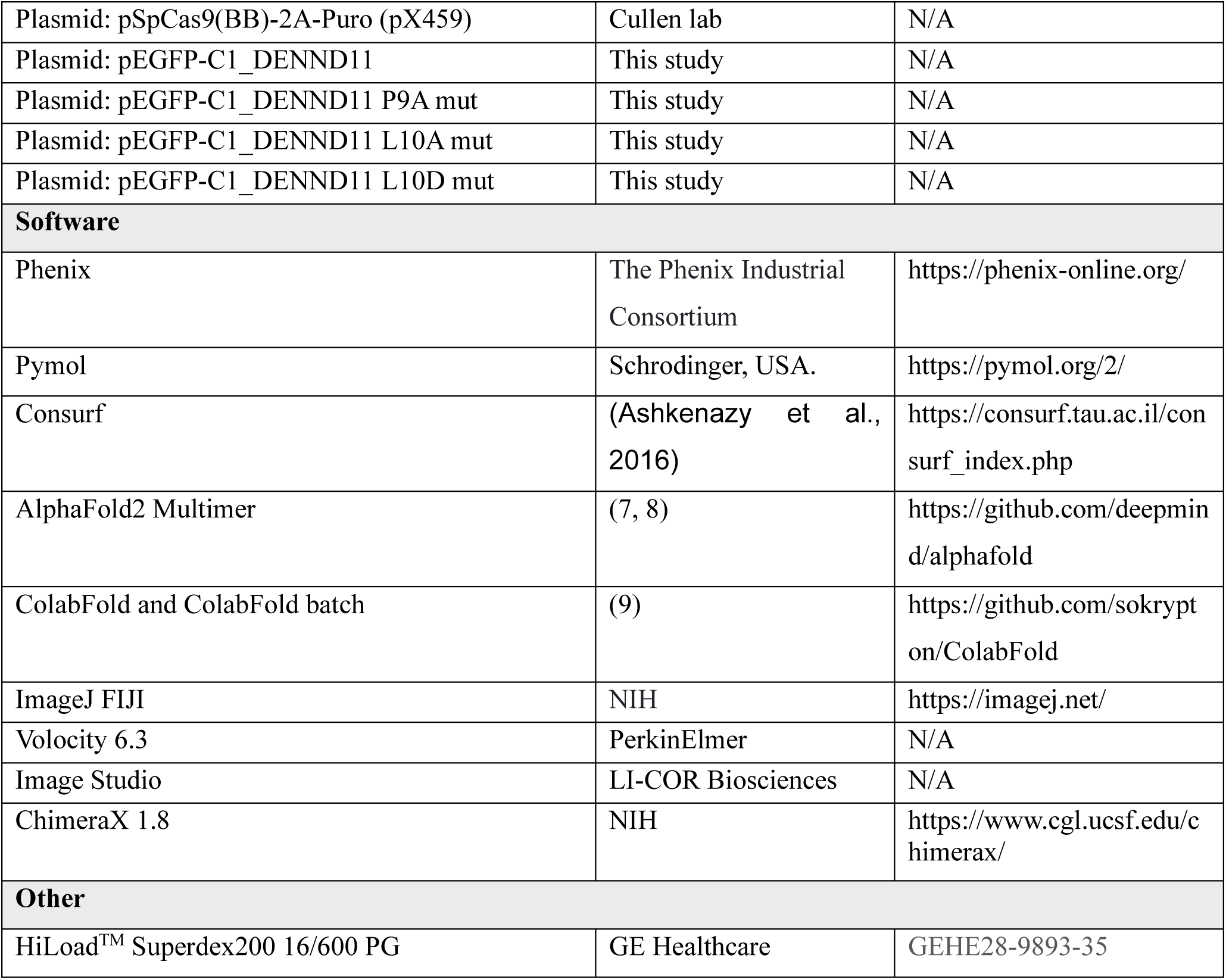
Reagent or Resource.

## Notes

### Competing Interest Statement

The authors have declared no competing interest.

